# Molecular underpinnings and biogeochemical consequences of enhanced diatom growth in a warming Southern Ocean

**DOI:** 10.1101/2020.07.01.177865

**Authors:** Loay Jabre, Andrew E. Allen, J. Scott P. McCain, John P. McCrow, Nancy Tenenbaum, Jenna L. Spackeen, Rachel E. Sipler, Beverley R. Green, Deborah A. Bronk, David A. Hutchins, Erin M. Bertrand

## Abstract

The Southern Ocean (SO) harbours some of the most intense phytoplankton blooms on Earth. Changes in temperature and iron availability are expected to alter the intensity of SO phytoplankton blooms, but little is known about how environmental change will influence community composition and downstream biogeochemical processes. We performed experimental manipulations on surface ocean microbial communities from McMurdo Sound in the Ross Sea, with and without iron addition, at −0.5 °C, 3 °C, and 6 °C. We then examined nutrient uptake patterns as well as the growth and molecular responses of two dominant diatoms, *Fragilariopsis* and *Pseudo-nitzschia*, to these conditions. We found that nitrate uptake and primary productivity were elevated at increased temperature in the absence of iron addition, and were even greater at high temperature with added iron. *Pseudo-nitzschia* became more abundant under increased temperature without added iron, while *Fragilariopsis* required additional iron to benefit from warming. We attribute the apparent advantage *Pseudo-nitzschia* shows under warming to upregulation of iron-conserving photosynthetic processes, utilization of iron-economic nitrogen assimilation mechanisms, and increased iron uptake and storage. These data identify important molecular and physiological differences between dominant diatom groups and add to the growing body of evidence for *Pseudo-nitzschia*’s increasingly important role in warming SO ecosystems. This study also suggests that temperature-driven shifts in SO phytoplankton assemblages may increase utilization of the vast pool of excess nutrients in iron-limited SO surface waters, and thereby influence global nutrient distributions and carbon cycle.

**Significance Statement:** Phytoplankton assemblages contribute to the Southern Ocean’s ability to absorb atmospheric CO_2_, form the base of marine food webs, and shape the global distribution of macronutrients. Anthropogenic climate change is altering the SO environment, yet we do not fully understand how resident phytoplankton will react to this change. By comparing the responses of two prominent SO diatom groups to changes in temperature and iron in a natural community, we find that one group, *Pseudo-nitzschia*, grows better under warmer low-iron conditions by managing cellular iron demand and efficiently increasing photosynthetic capacity. This ability to grow and draw down nutrients in the face of warming, regardless of iron availability, may have major implications for ocean ecosystems and global nutrient and carbon cycles.

## Main Text

The Southern Ocean (SO) occupies less than 10% of Earth’s ocean surface area but plays a major role in driving global climate and biogeochemical cycles. It connects the Pacific, Atlantic and Indian ocean basins, supplies nutrients to lower latitudes, and absorbs a considerable portion of global heat and CO_2_ (Frölicher et al. 2015; Moore et al. 2018). SO ecological processes are driven by seasonally productive phytoplankton assemblages that require iron and suitable temperatures to grow. These phytoplankton sequester atmospheric CO_2_ through the biological pump, sustain food webs, and affect dissolved nutrient availability globally. Climate change models predict overall warming trends in the SO (Turner et al. 2005; Ainley et al. 2010; Boyd et al. 2015; Moore et al. 2018; IPCC 2019) and changes in iron availability (Boyd et al. 2015; Hutchins and Boyd 2016; Tagliabue et al. 2017). Despite this, we still have a limited understanding of how these changes will influence the cellular mechanisms that govern phytoplankton dynamics and biogeochemistry in the region.

Low iron availability limits phytoplankton growth and is the primary cause of the high-(macro)nutrient low-chlorophyll (HNLC) conditions in the SO (de Baar et al. 1990; Martin et al. 1990). Despite this, diatoms are highly successful under these conditions, and contribute heavily to primary productivity throughout the SO (Arrigo et al. 1999; Kang et al. 2001). Low iron-adapted diatoms utilize several strategies to survive chronic, episodic, or seasonal iron limitation. For example, they can reduce photosynthetic iron demand by increasing the size of light harvesting antennae, and their photosynthetic electron transport chains (ETC) can substitute iron-containing ferredoxin with flavodoxin or cytochrome *c*_*6*_ with plastocyanin (Peers and Price 2006; Pankowski and McMinn 2009a; Strzepek et al. 2019). Diatoms may also use iron-free rhodopsin to supplement photosynthetic energy capture (Marchetti et al. 2015), have transport mechanisms that can be up-regulated to maximize iron acquisition (Allen et al. 2008; Morrissey et al. 2015; Kazamia et al. 2018; McQuaid et al. 2018; Coale et al. 2019), and contain iron storage proteins (ferritin) to buffer the sporadic availability of this micronutrient (Marchetti et al. 2009).

Temperature has also been shown to limit SO phytoplankton growth in the field and the laboratory. Shipboard warming incubations of iron-limited SO microbial communities increased nutrient drawdown and improved the growth of several diatom groups (Rose et al. 2009), with similar results reported in iron-limited laboratory cultures (Zhu et al. 2016). Warming increases the turnover rate of iron-containing enzymes including nitrate reductase (Gao et al. 2001; Di Martino Rigano et al. 2006), and may thus cause drawdown of SO nitrate even in the absence of iron addition (Hutchins and Boyd 2016; Spackeen et al. 2018). Increased temperature (up to a critical threshold) improves cellular iron use efficiency (Sunda and Huntsman 2011; Jiang et al. 2018) and may cause a reduction in the amount of enzymes -and iron-needed to carry out metabolic functions. Antarctic diatoms also exhibit reduced cell size under elevated temperature, which can increase surface area to volume ratios and facilitate nutrient uptake and growth (Zhu et al. 2016; Jabre and Bertrand 2020). Additionally, concurrent increases in temperature and iron have a much larger effect on growth compared to the individual effects of each factor (Rose et al. 2009; Zhu et al. 2016; Hutchins and Boyd 2016), with some species like *Fragilariopsis cylindrus* requiring iron supplementation to benefit from warming (Jabre and Bertrand 2020).

Despite shared adaptations across various taxa, diatoms are an extremely diverse group of phytoplankton. Within the SO, major diatom groups have different morphological and physiological traits, with unique optimal growth temperatures and different tolerances to low iron (Hutchins et al. 2001; Sackett et al. 2013; Tréguer et al. 2018; Strzepek et al. 2019). The molecular mechanisms that underpin these differences are still poorly understood, and are overlooked by the majority of current marine ecosystem models, which consider all diatoms as one group (Laufkötter et al. 2015). This oversimplification cannot capture the effects of specific diatom taxa on important ecological processes, or how environmentally mediated changes in diatom community composition and structure may affect nutrient and carbon biogeochemistry.

Here, we present an experimental manipulation study performed at the sea ice edge in the Ross Sea of the Southern Ocean that investigated the growth, nutrient drawdown, and molecular responses of a SO microbial community to warming and changes in iron availability. We show that nutrient drawdown increases in response to temperature and iron, both independently and synergistically, and that two closely related diatom groups, *Fragilariopsis* spp. and *Pseudo-nitzschia* spp., respond differently to changes in iron and temperature. These two taxa contribute substantially to diatom assemblages in the SO, and strongly influence the ecology and biogeochemistry of the region (Kang and Fryxell 1992; Rose et al. 2009). *Pseudo-nitzschia* cell numbers increased more than *Fragilariopsis* under warming alone, while the latter required iron addition to benefit from increased temperature. Metatranscriptomes suggest that growth of *Pseudo-nitzschia* under warming is likely facilitated by temperature-responsive light harvesting and iron management strategies that were not utilized by *Fragilariopsis*. This ability to grow and draw down nutrients in the face of warming, regardless of iron availability, could have major implications for the ecological and biogeochemical response of the HNLC SO to a changing climate.

## Results and Discussion

We incubated SO surface water with and without iron addition at −0.5 °C, 3°C, and 6 °C, and examined the microbial community after 24 hours (T1) and again after five (T5) and seven days (T7) (Methods). Increasing temperature and iron separately caused a significant but small increase in primary productivity measured as bicarbonate uptake (Fig. 1A; Fig. S1, two-way ANOVA, temperature effect *p* = 1.1 ×10^−6^, Fe effect *p* = 2.4 × 10^−6^), as well as an increase in nitrate uptake rates (Fig.1A; Fig. S1, two-way ANOVA, temperature effect *p* = 0.001, Fe effect *p* = 7.9 ×10^−5^), nitrate consumption (Fig. 1B; Fig. S1, two-way ANOVA, temperature effect *p* = 2.9 ×10^−11^, Fe effect *p* = 1.2 ×10^−11^) and phosphorus consumption (Fig. 1B, Fig. S1, two-way ANOVA, temperature effect *p* = 2.3 ×10^−6^, Fe effect *p* = 1.0 ×10^−6^). Incubations under concurrent warming and iron addition showed an even larger increase in primary productivity (Fig. 1A, two-way ANOVA, temperature x Fe effect *p* = 4.7 ×10^−5^), nitrate uptake rates (Fig. 1A, two-way ANOVA, temperature x Fe effect *p* = 0.007), nitrate consumption (Fig. 1B; Fig. S1, two-way ANOVA, temperature x Fe effect *p* = 6.0 ×10^−9^), and phosphorus consumption (Fig. 1B, Fig. S1, two-way ANOVA, temperature x Fe effect *p* = 7.5 × 10^−5^). This synergistic iron-temperature effect has been observed previously (Rose et al. 2009; Spackeen et al. 2018), and shows that in addition to iron, temperature plays an important role in constraining SO phytoplankton growth and macronutrient utilization. These data suggest that future warming conditions can strongly influence SO productivity and nutrient drawdown, especially under increased iron availability.

**Figure 1.**
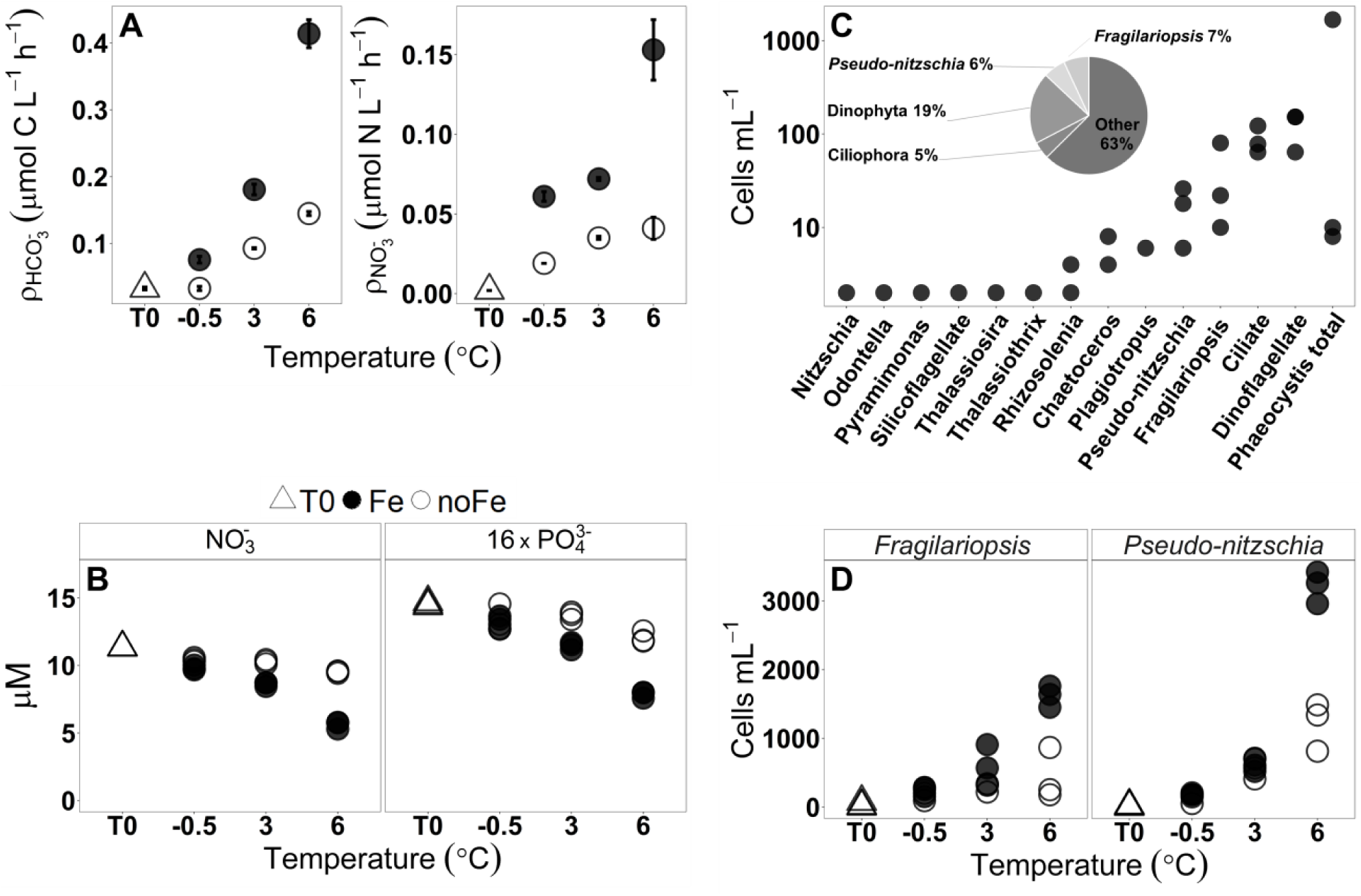
**A)** Absolute bicarbonate and nitrate uptake rates at T0 (prior to the incubations) and T7 for each temperature and iron condition. Uptake rates were measured in duplicate from a single incubation bottle per treatment. **B)** Dissolved nitrate and phosphate concentrations in triplicate incubation bottles at T0 and T7. Phosphate concentration is scaled (16 x) for better visual comparison with nitrate. **C)** Triplicate measurements of initial (T0) cell counts of eukaryotic taxa identified by light microscopy. Taxa are arranged from low to high abundance. Inset pie chart represents percent contribution of taxa to the total reads mapped to ORFs. “Other” represents all other taxonomic groups, including prokaryotes and viruses. **D)** Measurements of *Fragilariopsis* and *Pseudo-nitzschia* cell counts at T0 and T7 in triplicate incubation bottles for each temperature and iron condition.

Microscopy showed that *Phaeocystis*, dinoflagellates, ciliates, *Fragilariopsis* and *Pseudo-nitzschia* were among the most abundant eukaryotic taxa at the beginning of the incubation experiment (the *in-situ* community), and after temperature and iron incubations (Fig. 1C; Fig. S2). We focused our analyses here on the responses of *Fragilariopsis* (mostly *F. kerguelensis*) and *Pseudo-nitzschia* (mostly *P. subcurvata)*, which play integral roles in SO ecology. *Pseudo-nitzschia* but not *Fragilariopsis* cell counts increased significantly under warming without added iron (Fig. 1D, two-way ANOVA, temperature effect *p* = 0.003 and *p* = 0.5 respectively). Additionally, concurrent warming and iron supplementation caused an increase in *Fragilariopsis* (Fig. 1D, two-way ANOVA, temperature x iron effect *p* = 0.001), and a much larger increase in *Pseudo-nitzschia* cell counts (Fig. 1D, two-way ANOVA, temperature x iron effect *p* = 1.13 × 10^−6^. This highlights the ability of *Pseudo-nitzschia* to take advantage of increased temperature under both low and high iron availability.

We used metatranscriptomics to comprehensively capture gene expression patterns and examine the molecular processes underpinning the distinct growth responses of *Fragilariopsis* and *Pseudo-nitzschia* to shifts in iron availability and temperature (Fig. 2). Metatranscriptomics allows for the interrogation of cellular pathways utilized across various taxa, and can be used to infer metabolic states under different environmental conditions (Marchetti et al. 2012; Bertrand et al. 2015; Cohen et al. 2018a). Here we identified transcripts belonging to viruses, prokaryotes, and eukaryotes (Fig. S3); with dinoflagellates, ciliates, *Pseudo-nitzschia* and *Fragilariopsis* contributing 37% of the total reads mapped to ORFs (Fig. 1C). The contribution of *Phaeocystis* to the metatranscriptome was lower than anticipated based on microscopy, possibly due to difficulty in harvesting cells/colonies without rupturing them (Kiene and Slezak 2006). KEGG Orthology (K.O.) term enrichment analysis on ORFs belonging to *Fragilariopsis* and *Pseudo-nitzschia* showed that pathways corresponding to photosynthesis, nitrogen metabolism and genetic information processing/translation were highly differentially expressed in both taxa following temperature and iron increase (Fig. S4; Fig. S5). We explored these pathways in more detail by grouping individual ORFs into groups of similar sequences (clusters) using Markov clustering (MCL) (Enright 2002) (Methods). Several clusters contained differential expression patterns that were similar in *Fragilariopsis* and *Pseudo-nitzschia* under iron addition (63 clusters) and temperature increase (212 clusters). However, a larger number of clusters were uniquely differentially expressed in either *Fragilariopsis* or *Pseudo-nitzschia* under these conditions (Fig. 3A), suggesting that *Fragilariopsis* and *Pseudo-nitzschia* utilize different molecular strategies to respond to changes in temperature and iron.

**Figure 2.**
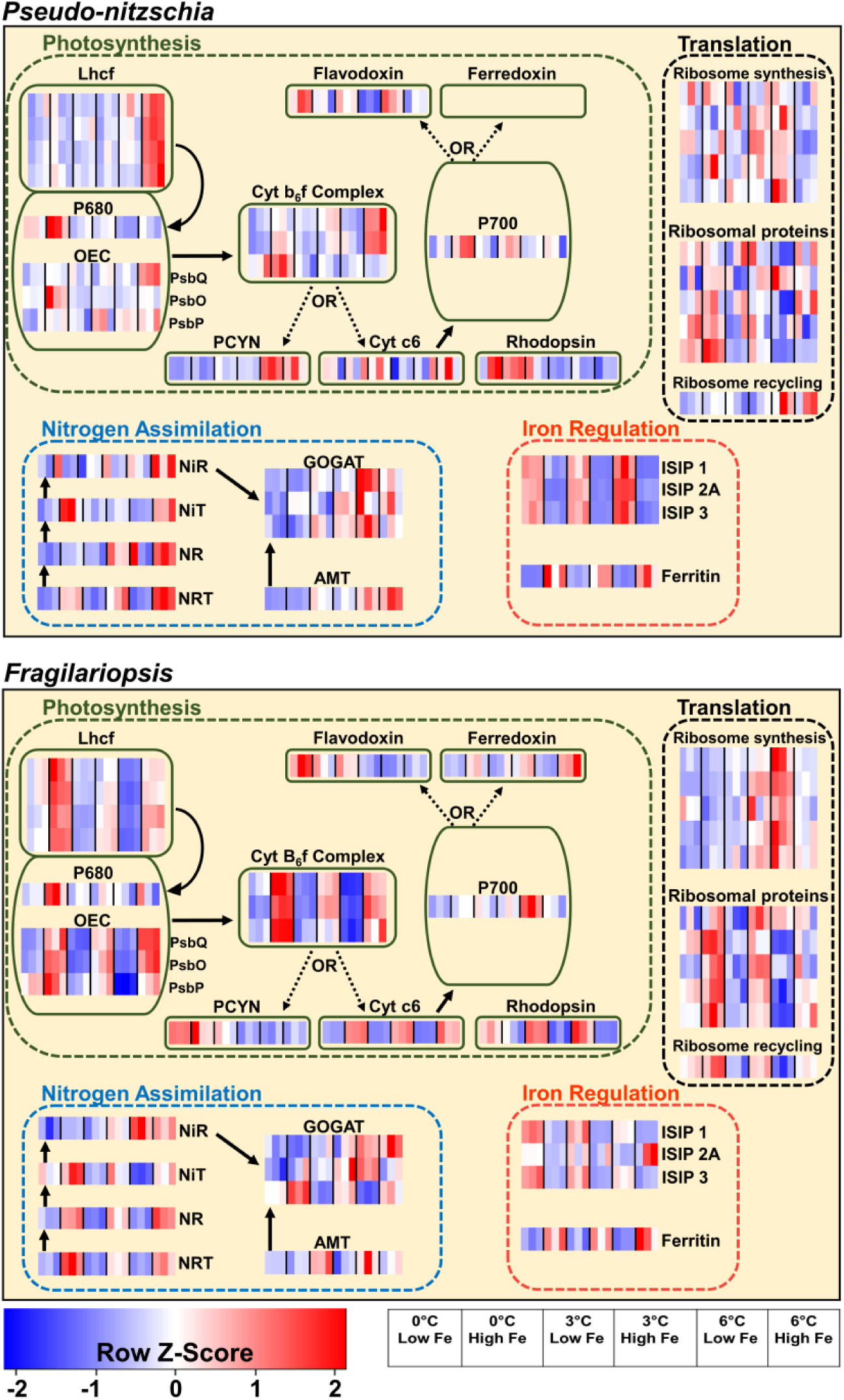
Schematic representations of *Pseudo-nitzschia* and *Fragilariopsis* cells showing cellular processes, with each process comprised of several protein clusters (MCL clusters). Lhcf = light harvesting complexes-f, OEC = oxygen evolving complex, Cyt B6f complex = cytochrome b_6_f complex, PCYN = plastocyanin, Cyt c6 = cytochrome c6, NRT = nitrate transporter, NR = nitrate reductase, NiT = nitrite transporter, NiR = nitrite reductase, AMT = ammonium transporter, GOGAT = glutamine oxoglutarate aminotransferase cycle, ISIP = iron starvation induced protein. Each row in a heatmap represents one Markov cluster (MCL), each column represents a temperature and iron treatment at T5 with each block representing one biological replicate. Heatmaps were constructed using taxon-normalized RPKM values Empty heatmap placeholders represent clusters found in *Fragilariopsis* but not *Pseudo-nitzschia*. Arrows represent energy/electron flow in photosynthetic light reactions, and steps involved in nitrogen assimilation using nitrate or ammonium.

**Figure 3.**
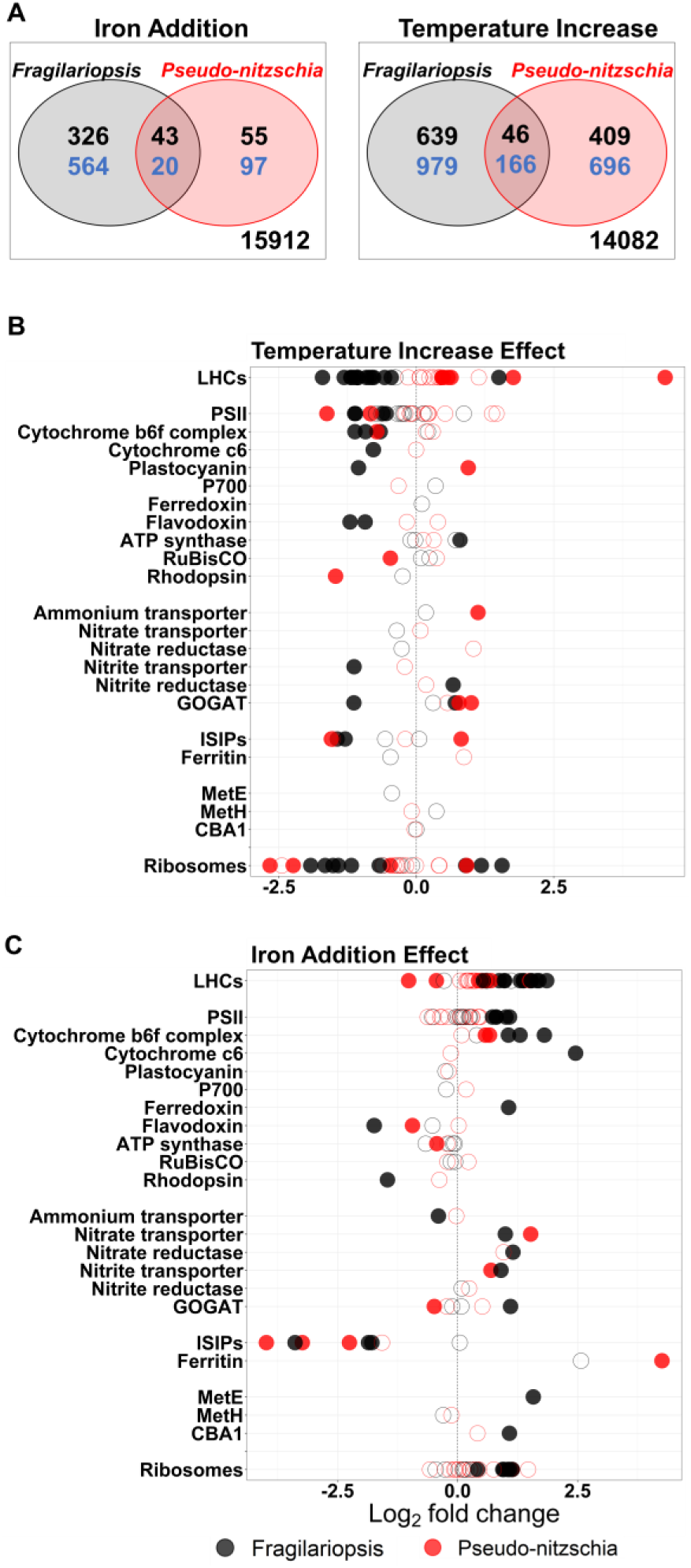
**A)** Venn diagram showing the number of transcript clusters that are significantly up (black) or down (blue) regulated with iron addition or temperature increase in *Fragilariopsis* and *Pseudo-nitzschia*. The intersection represents clusters up or down regulated in both *Fragilariopsis* and *Pseudo-nitzschia.* The number outside of the circles represents the number of clusters from both taxa that were not differentially expressed. In both taxa, warming caused more clusters to be differentially expressed than iron addition. **B)** Differential expression (DE) of various clusters under increased temperature. Differential expression was calculated using the quasi likelihood test (glmQLFTest) in EdgeR, and fold-change was calculated for high-temperature (6 °C) vs low temperature (−0.5 °C) treatments at T5. **C)** Differential expression (DE) of various clusters after iron addition. Differential expression was calculated using the quasi likelihood test (glmQLFTest) in EdgeR and fold-change was calculated for iron-added vs no iron-added treatments at T5. In both B and C, positive and negative Log2 foldchange values represent up and down regulation respectively. Filled circles are clusters with statistically significant DE (adjusted p-value <0.05).

### Photosynthesis

Iron addition resulted in upregulation of iron-containing cytochrome b_6_f complex transcripts and downregulation of flavodoxin transcripts in both *Fragilariopsis* and *Pseudo-nitzschia* (Fig. 2, Fig. 3). This indicates that both diatoms in our study were experiencing some degree of iron stress at the time of sampling, as iron-containing photosynthetic processes including the electron transport chain can account for up to 40% of cellular iron quotas (Raven et al. 1999) and are typically downregulated under low iron (LaRoche et al. 1996; Allen et al. 2008). Here we did not detect the expression of iron-dependent electron acceptor ferredoxin (PetF) in *Pseudo-nitzschia*, regardless of iron status. Antarctic *Pseudo-nitzschia* sp. may have constitutively reduced dependence on petF (Pankowski and McMinn 2009b; Moreno et al. 2018) to minimize photosynthetic iron demand.

### Light Harvesting Complexes

Most members of the light-harvesting complex (LHC) superfamily primarily absorb and direct light energy to photosynthetic reaction centers where charge separation occurs, but some of them (Lhcx clade) are also involved in stress responses and photoprotection (Zhu and Green 2010; Kirilovsky and Büchel 2019; Buck et al. 2019). Expression patterns of transcripts belonging to the major Lhcf group and PSI-associated Lhcr groups were markedly different between the two taxa (Fig. 4). In *Fragilariopsis*, there was a general pattern of decreased expression with warming, and marked upregulation with iron addition. In contrast, *Pseudo-nitzchia* showed a pattern of increasing expression with elevated temperatures. Upregulation of Lhcf and Lhcr groups contributes to increased light harvesting efficiency, and alleviates photosynthetic iron demand by reducing the number of iron-containing photosynthetic components required to process light energy (Schuback et al. 2015). Increased light harvesting cross section under warming has been detected in iron-stressed *F. cylindrus* under laboratory conditions (Jabre and Bertrand, 2020), and these environmental data indicate that *Pseudo-nitzschia* may be even better equipped to use this mechanism to support growth under low iron in the SO. Several Lhcf and Lhcr clusters were also upregulated in both *Fragilariopsis* and *Pseudo-nitzschia* after iron addition. Given that iron addition is expected to reduce light harvesting cross section (Greene et al. 1992; Jabre and Bertrand 2020) it might be expected that Fe addition could decrease demand for Lhcf and Lhcr expression. However, this impact appears to be overridden by increased demand for LHC expression as a result of Fe-induced increases in PSU abundance. Additional photophysiological measurements could provide further insight into how light harvesting cross section responds to changes in LHC expression under different iron and temperature conditions.

**Figure 4.**
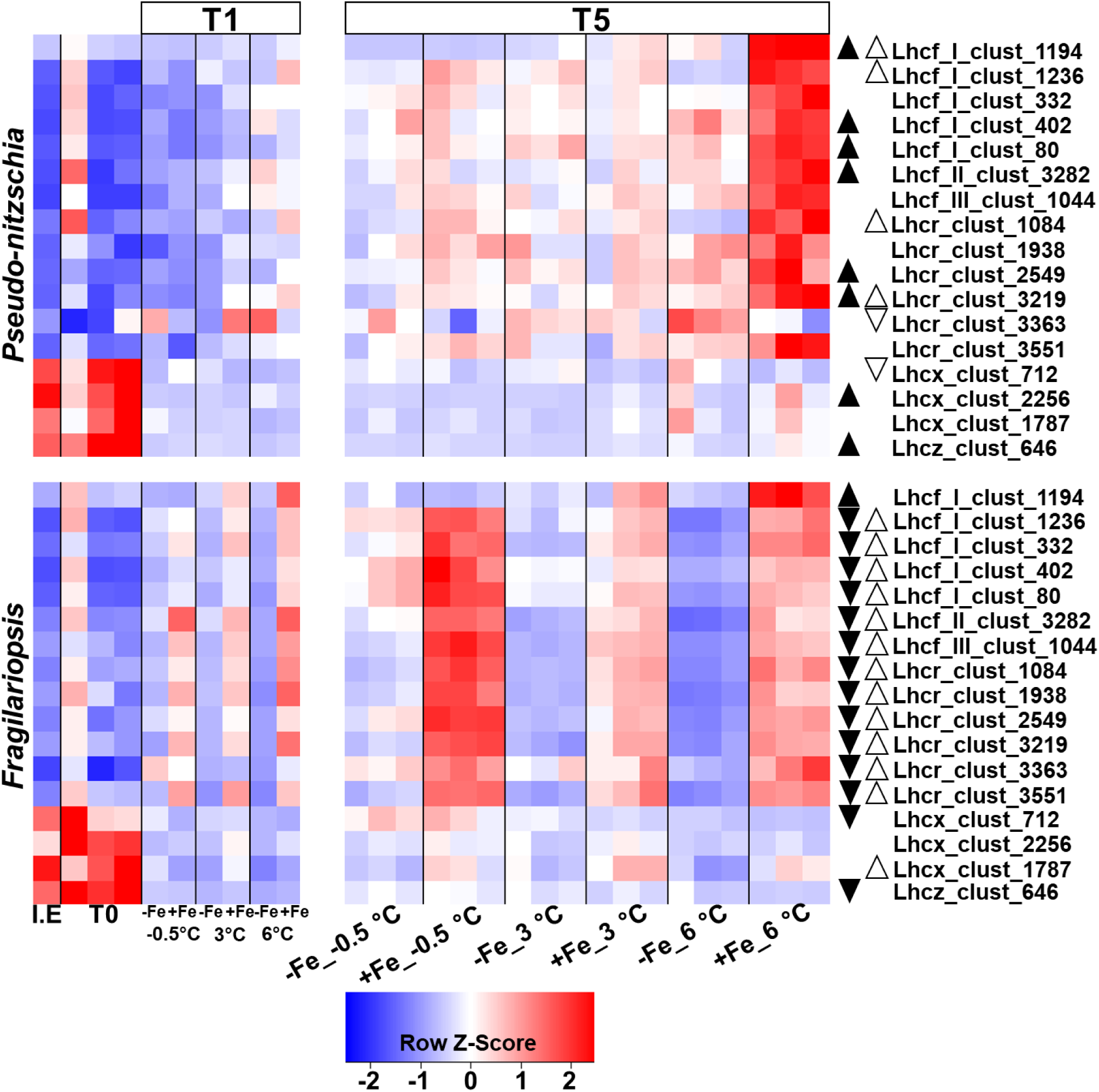
Heatmaps of MCL clusters of light harvesting complexes Lhcf, Lhcr, Lhcx, Lhcz (see Table S3) in *Pseudo-nitzschia* and *Fragilariopsis* measured after 24 hours (T1) and 5 days (T5) of incubation under the various iron and temperature treatments. Heatmaps were constructed using taxon-normalized RPKM values. I.E represents ice edge samples, T0 represents *in-situ* samples processed immediately before any incubations. Each block is one biological replicate measurement. White-filled up/down pointing triangles represent transcripts that were significantly (glmQLFTest-EdgeR p <0.05) up or down regulated due to iron addition at T5. Black-filled up/down pointing triangles represent transcripts that were significantly (glmQLFTest-EdgeR p <0.05) up or down regulated due to warming at T5.

Notably, each transcript cluster belonging to the Lhcx clade was downregulated after 24 hours of incubation at all temperatures and iron conditions in both *Fragilariopsis* and *Pseudo-nitzchia* (Fig. 4). Lhcxs are highly sensitive to environmental change, including fluctuations in light and nutrient levels (Zhu and Green 2010; Taddei et al. 2016; Buck et al. 2019). Given these expression patterns and previous observations of Lhcx regulation, the Lhcx transcripts detected here may be responding to changes in light levels during sample collection. Notably, expression of these Lhcx clades showed only minimal responses to iron addition.

### Plastocyanin

Plastocyanin is a copper-containing protein that acts as an electron shuttle in the ETC and plays an important role in reducing iron requirements in phytoplankton by substituting for cytochrome c_6_ (Peers and Price 2006; Raven 2013; Wu et al. 2019). Here, iron addition had no significant effect on plastocyanin transcript expression, while warming caused its upregulation in *Pseudo-nitzschia* and downregulation in *Fragilariopsis* (Fig. 3). Plastocyanin expression has been observed to be insensitive to iron availability in low iron adapted diatoms (Marchetti et al. 2012). However, our data suggest that *Pseudo-nitzschia* plastocyanin transcripts are phylogenetically distinct from those of *Fragilariopsis*, and may play a temperature-responsive role to support growth under elevated temperatures (Fig. S6). Plastocyanin transcript abundance in *Pseudo-nitzschia* was higher at 3 °C and 6 °C compared to −0.5 °C at T1, showing a rapid response to warming (Fig. S6). This, in combination with the strong and immediate response of LHCs may give *Pseudo-nitzschia* a photosynthetic advantage over *Fragilariopsis* in warmer waters.

### Nitrogen Metabolism

K.O. term enrichment analysis showed that transcripts encoding nitrogen metabolism were enriched in *Pseudo-nitzschia* and depleted in *Fragilariopsis* after temperature increase (Fig. S4; Fig. S5). Nitrogen is required for protein synthesis and would be necessary to support the observed increase in *Pseudo-nitzschia* cell counts under warming, and notably, dissolved nitrogen drawdown was enhanced due to warming alone and elevated further upon iron addition (Fig. 5A, B). However, nitrogen acquisition and assimilation from nitrate requires iron (Schoffman et al. 2016) and the observed *Pseudo-nitzschia* growth under elevated temperature would necessitate iron-economic nitrogen metabolism. Our data show that ammonium transporter transcripts were upregulated in *Pseudo-nitzschia* and not in *Fragilariopsis* under warming (Fig. 3C; Fig. 5). Previous work with laboratory cultures suggests that ammonium transporter expression is regulated by nitrogen demand and can be independent of ammonium availability (Bender et al. 2014). This indicates that *Pseudo-nitzschia* utilizes ammonium transporters in response to elevated nitrogen demand, which is likely a result of increased growth in response to warming. Elevated ammonium transporter expression would provide *Pseudo-nitzschia* with an additional nitrogen source at a lower iron cost for use in amino acid synthesis and growth (Raven 1988; Schoffman et al. 2016; Smith et al. 2019). In fact, ammonium drawdown at 6 °C was greater than at lower temperatures, regardless of iron status (Fig. 5B), consistent with the notion that ammonium use at high temperature and low iron could have contributed to the success of *Pseudo-nitzschia*. However, the high *Pseudo-nitzchia* growth we observed cannot be supported by ammonium alone; ammonium drawdown comprised only 25.2 ± 2.4 % of dissolved inorganic nitrogen (nitrate + ammonium) drawdown in the 6° no added iron treatment and 7.9 ± 1.0 % of dissolved inorganic nitrogen drawdown in the 6° with added iron treatment (Fig. 5A, B). Nitrate drawdown was responsible for the majority of dissolved nitrogen uptake regardless of iron status, though significantly (two-way ANOVA, *p* = 9.6 ×10^−11^) more so under elevated iron. While nitrate drawdown was elevated at high temperatures both with and without added iron (Fig. 5A), neither *Pseudo-nitzschia* nor *Fragilariopsis* changed their expression of nitrate transporters in response to elevated temperature. Both diatom groups up-regulated nitrate transporters in response to Fe addition regardless of temperature (Fig. 3; Fig. 5E, F). Similarly, nitrate reductase transcript expression was up-regulated in response to Fe addition but not elevated temperature (Fig. 3; Fig. 5G, H). However, the observed elevation in nitrate drawdown and nitrate uptake rates with warming (Fig. 1; Fig. 5), in the absence of a change in gene expression, is consistent with the well-characterized temperature dependence of nitrate reductase activity (Gao et al. 2001). This suggests that the enhanced nitrate assimilation under elevated temperature was indeed accomplished without the increase in iron demand required to produce additional copies of active nitrate reductase.

**Figure 5.**
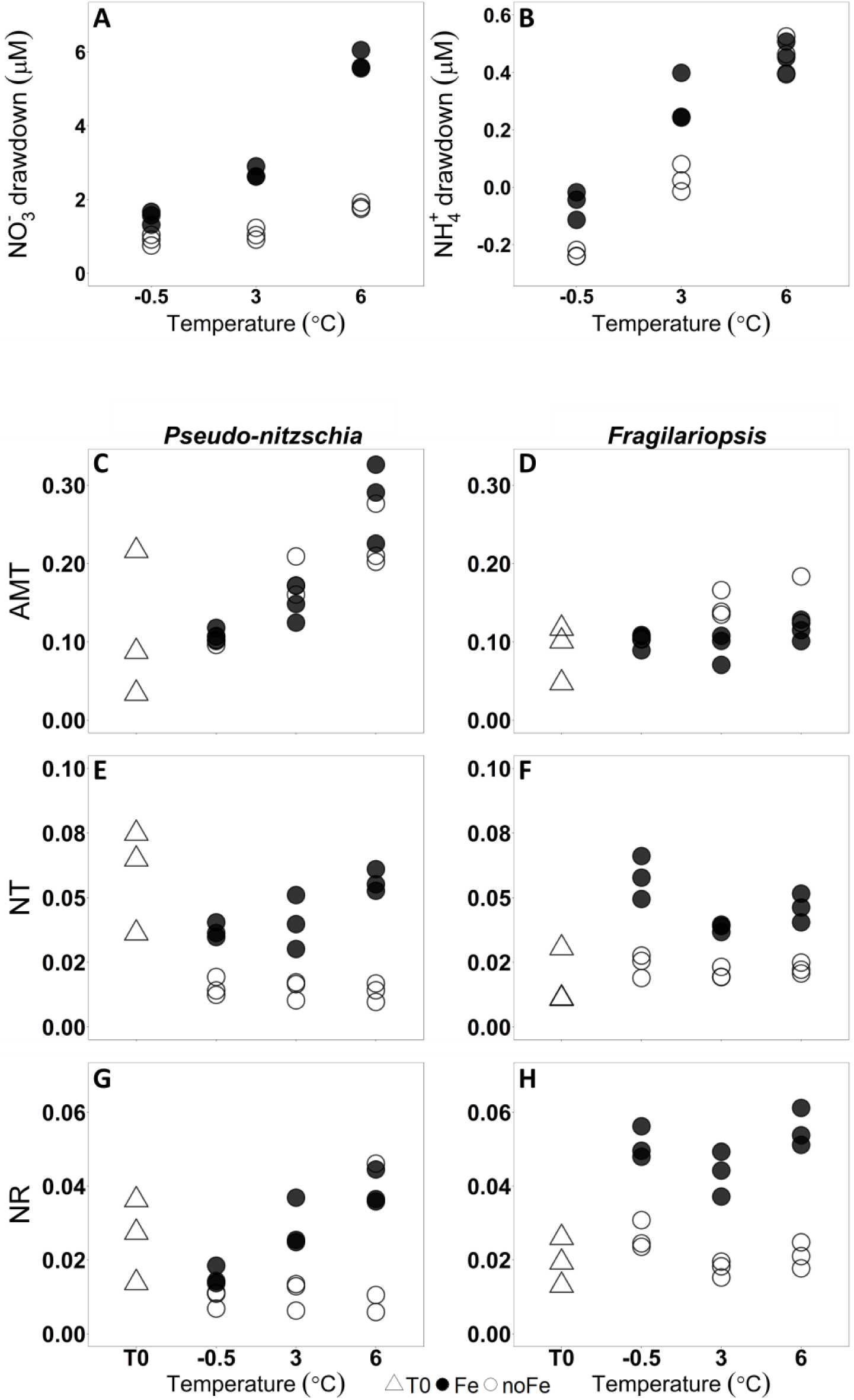
**A, B)** Total nitrate and ammonium drawdown from beginning to end of the experiment (difference in concentration between T7 and T0) **C-H)** Taxon normalized expression values for clusters belonging to ammonium transporters (AMT) nitrate transporters (NT) and nitrate reductase (NR) in both *Pseudo-nitzschia* and *Fragilariopsis* prior to incubation (T0), and at T5 with and without added iron at −0.5, 3, and 6 °C.

### Genetic Information Processing / Translation

Protein synthesis is an energetically costly process that is required for metabolism and growth. Increased kinetic energy due to warming enhances ribosomal efficiency, and reduces the number of ribosomes required to maintain protein synthesis rates (Toseland et al. 2013). Consistent with this response, we observed that several transcript clusters corresponding to ribosomes were downregulated in both *Fragilariopsis* and *Pseudo-nitzschia* under warming (Fig. 2, Fig. 3). Warming also caused upregulation of transcripts encoding ribosomal recycling proteins only in *Pseudo-nitzschia* (Fig. 2), which can reduce the energetic costs associated with ribosomal synthesis. The reduction in phosphorus-rich ribosome abundance increases N:P ratios in warmer ocean regions (Toseland et al. 2013). However, bulk nutrient drawdown ratios measured here (Fig. S1) show that N:P drawdown is not influenced by warming (two-way ANOVA, *p* = 0.07). This suggests that the increasing dominance of diatoms, with their considerably lower N:P ratios than other SO plankton groups (Arrigo et al. 1999; Zhu et al. 2016), is likely an overall driver of N:P and can counterbalance the expected reduction in cellular phosphate demand at high temperature from reduced ribosome content.

### Cobalamin Metabolism

Exogenous vitamin B_12_ (cobalamin) acts as a cofactor in the cobalamin-requiring methionine synthase enzyme (MetH) found in all diatoms to synthesize the essential amino acid methionine and facilitate one-carbon metabolism (Bertrand and Allen 2012). Under low cobalamin availability, however, certain diatoms including *Fragilariopsis* but not *Pseudo-nitzschia*, are capable of synthesizing methionine from homocysteine using a less efficient cobalamin-independent methionine synthase (MetE) (Bertrand et al. 2012; Ellis et al. 2017). Diatoms also upregulate CBA1, a cobalamin acquisition related protein, under cobalamin deprivation (Bertrand et al. 2012; Ellis et al. 2017). Following the initial 24-hour incubation period, CBA1 and MetH were upregulated in *Pseudo-nitzschia* after warming at 6 °C (Fig. S7), but were not significantly influenced by iron or temperature following 5-day incubations (Fig. 3; S7). In contrast, MetE and CBA1 transcripts were significantly upregulated in *Fragilariopsis* following iron additions after 5-day incubations (Fig. 3; S7), suggesting rearranged metabolism to cope with cobalamin deprivation that likely emerged in response to iron addition (Bertrand et al. 2015).

Rapid cobalamin uptake by *Pseudo-nitzschia*, facilitated by upregulation of CBA1 after 24 hours, may give it a competitive advantage when cobalamin is available. However, the ability of *Pseudo-nitzschia* to maintain vigorous growth without significantly elevating CBA1 expression after 5 days is notable, and suggests that these two diatoms may employ different strategies to cope with low cobalamin availability. *Fragilariopsis* appears to reduce cobalamin demand through the use of MetE while increasing investment in acquisition with CBA1. In contrast, *Pseudo-nitzschia* may 1) have a MetH enzyme that is more efficient at elevated temperatures compared to *Fragilariopsis*, 2) employ salvage and repair of degraded cobalamin complexes (Cohen et al. 2018a) or 3) rely on bacteria in close physical association for cobalamin supply. Our data show that the overall expression of genes encoding cobalamin salvage and remodeling proteins were not influenced by iron status or temperature in *Pseudo-nitzschia* (Fig. S7), despite previous evidence of their upregulation with iron addition in North Pacific diatoms including *P. granii* (Cohen et al. 2017, 2018a). Further work comparing cobalamin uptake and methionine synthase kinetics between *Pseudo-nitzschia* and *Fragilariopsis*, and examining their relationships with cobalamin producing and consuming bacteria, could provide further insight into how these taxa cope with episodic decreases in cobalamin availability.

### Iron acquisition, trafficking, and storage

Increased iron uptake, storage, and intracellular transport are strategies to improve growth under low iron availability. Zhu et al. (2016) report iron uptake rates and Fe:C ratios that are higher in *Pseudo-nitzschia* under increased temperature, and no warming effect on iron acquisition in *Fragilariopsis*. Our data show downregulation of iron stress induced proteins (ISIP1, ISIP2A, ISIP3) in both *Pseudo-nitzschia* and *Fragilariopsis* after iron addition, and ISIP2A upregulation only in *Pseudo-nitzschia* after warming (Fig. 2; Fig. 3). ISIPs are involved in iron acquisition and storage systems in diatoms, and specifically, ISIP2A contributes to non-reductive iron uptake (Allen et al. 2008; Morrissey et al. 2015; Kazamia et al. 2018; McQuaid et al. 2018). No previous studies have inspected ISIP2A response to temperature, but it is likely that the apparent temperature responsiveness of ISIP2A contributes to the ability of *Pseudo-nitzschia* to acquire and utilize iron to support growth under warming. Additionally, transcripts for ferritin, an iron storage protein (Marchetti et al. 2009), were significantly upregulated only in *Pseudo-nitzschia* after iron addition (Fig. 3). Several studies have reported ferritin upregulation in *Pseudo-nitzschia* under iron replete conditions (Cohen et al. 2018b; a), and *Fragilariopsis* is also believed to utilize ferritin for long-term iron storage (Marchetti et al. 2009). However, similar to what has been observed in previous field studies (Lampe et al. 2018), *Fragilariopsis* ferritin was not significantly upregulated after iron addition here (Fig. 3). These trends suggest that in addition to reducing iron demand, *Pseudo-nitzschia* was better able to increase labile iron uptake and utilized stored ferritin-bound iron to support growth under warming.

### Ecological and biogeochemical relevance

Most recent model projections show increased temperature trends across the SO with slight warming in areas previously thought not likely to be affected by climate change (Boyd et al. 2015; Rickard and Behrens 2016; Moore et al. 2018). *Fragilariopsis* grows in a wide range of environments throughout the SO but is most successful in high-salinity, high-iron sea ice (∼ 0 °C), while *Pseudo-nitzschia* is most successful in low-salinity, low-iron meltwaters (∼ 2 °C) (Petrou et al. 2011; Petrou and Ralph 2011; Sackett et al. 2013). The growth and gene expression patterns we observed in both diatoms may be driven by niche-specific adaptations due to competition for resources. Our results are consistent with a 2013 SO incubation study where *Pseudo-nitzchia* dominated the community after warming (Spackeen et al. 2018), and suggest that Antarctic *Pseudo-nitzschia* are better equipped to dominate in a warmer ocean, and even more so if iron availability increases. A shift into a *Pseudo-nitzschia* dominated community raises concerns for domoic acid (DA) production and its toxic effects (Bates et al. 1998). A search for DA biosynthesis transcripts (Brunson et al. 2018) did not yield strong evidence for expression of the DA biosynthesis gene cluster in this experiment (Table S1; Table S2). However, a few studies have reported DA production in the SO (Silver et al. 2010; Geuer et al. 2019), but the temporal, spatial and environmental factors that elicit DA production remain unclear. Further work is required to investigate whether resident *Pseudo-nitzschia* can produce the toxin under conditions not examined in our study, or if DA producing *Pseudo-nitzschia* are likely to migrate into a warmer SO.

A change in SO phytoplankton community structure could have important biogeochemical ramifications on local and global scales. Notably, warming-induced increases in iron-conserving plastocyanin, LHCs and nitrogen assimilation mechanisms observed here likely resulted in improved iron use efficiency (lower Fe:C uptake ratios) (Zhu et al. 2016). Elevated iron use efficiency with warming appears to result in enhanced nutrient drawdown even in the face of iron limitation. Such warming-enhanced SO nutrient drawdown could have profound consequences for global nutrient distribution. As more nutrients are consumed by phytoplankton and exported out of the surface ocean around Antarctica, less nutrients are available to fuel productivity at lower latitudes (Moore et al. 2018). Here we have identified molecular mechanisms that enable diatoms, particularly *Pseudo-nitzschia*, to accomplish enhanced growth and nutrient drawdown with rising temperatures under low iron conditions. These mechanisms; upregulation of iron-conserving photosynthetic processes, utilization of iron-economic nitrogen assimilation mechanisms, and increased iron uptake and storage; underpin what may be a pattern of increasing nutrient utilization in the Southern Ocean and decreasing availability of nutrients for high latitudes.

## Materials and Methods Summary

Seawater was collected at a 3-meter depth at the ice edge from McMurdo Sound, Antarctica on January 15^th^, 2015 (165°24.7985’E, 77°37.1370’S) using previously described trace-metal-clean techniques (Bertrand et al. 2015). Triplicate bottles of each treatment (temperature: −0.5 °C ± 0.2 °C, 3 ± 0.5 °C and 6 ± 0.5°C, iron (2 nM): not added, added) were incubated indoors at a constant irradiance of 65-85 uE m^-2^ sec^-1^. Metatranscriptome samples were taken at the sea ice edge (IE), in the laboratory immediately following bottle incubation setup (T0), on January 16^th^ (T1), and again on January 20^th^ (T5). Cell counts were measured on T0 and January 22^nd^ (T7). Dissolved nutrients (nitrate, phosphate, silicate) were measured on T0, T1, T3, T5 and T7, and ammonium was measured on T0 and T7. Primary productivity (bicarbonate uptake) and nitrate uptake rates were measured on T0, T1, T3 and T7 (Fig. S1).

For metatranscriptomics, the microbial community was harvested on 0.2 μm Sterivex™ filters, total RNA was extracted using Trizol reagent (Thermo Fisher Scientific) and ribosomal RNA was removed with Ribo-Zero Magnetic kits (Illumina). The resulting rRNA was purified and subjected to amplification and cDNA synthesis, using the Ovation RNA-Seq System V2 (TECAN). One microgram of the resulting cDNA pool was fragmented to a mean length of 200 bp, and libraries were prepared using Truseq kit (Illumina) and subjected to paired-end sequencing via Illumina HiSeq. Illumina paired reads were filtered to eliminate primer sequences and quality trimmed to Phred Q33, and rRNA identified and removed using riboPicker (Schmieder et al. 2012). Transcript contigs were assembled de novo using CLC Assembly Cell (http://www.clcbio.com) and open reading frames (ORFs) were predicted using FragGeneScan (Rho et al. 2010). ORFs were annotated for putative function using hidden Markov models and blastp against PhyloDB (Bertrand et al. 2015), and filtered to eliminate those with low mapping coverage (< 50 reads total over all samples), proteins with no blast hits and no known domains. ORFs were assigned to taxonomic groups of interest (Fig. S2) based on best LPI taxonomy (Podell and Gaasterland 2007; Bertrand et al. 2015). Reads per kilobase mapped (RPKM) expression values for each ORF were calculated and taxon normalized using a normalization factor representing the summed taxonomic group contribution to total nuclear-assigned reads per sample. ORFs were clustered into orthologs and protein families using MCL (Enright 2002). Group normalized cluster (average/total) RPKM expression values were calculated by pooling the taxon-normalized expression values for each group within a cluster. Cluster annotations were aggregated by annotation type (KEGG, KO, KOG, KOG class, Pfam, TIGRfam, EC, GO) and a single annotation was chosen to represent each cluster based on the lowest Fisher’s exact test p-value (fisher.test in R).

Differential gene expression at T5 (foldchange magnitude and adjusted p-value) was calculated using empirical Bayes quasi-likelihood F-tests (glmQLFTest in edgeR) with a p-value cut off of 0.05 for statistical significance. Iron effect was determined by comparing all -Fe treatments at T5 to all +Fe treatmenets at T5. Temperature effect was determined by comparing all −0.5 °C to 3 °C treatments at T5, and −0.5 °C to 6 °C at T5. A gene was considered upregulated with temperature if it was upregulated in both −0.5 °C vs 3 °C and −0.5 °C vs 6 °C tests.

Stable isotope tracer techniques, using ^15^N-labeled nitrate and ^13^C-labeled bicarbonate (Cambridge Isotope Laboratories), were used to determine uptake rates, similar to methods previously described in Ross Sea nutrient utilization studies (Spackeen et al. 2018). Tracer amounts of labeled substrates were added to bottles containing the initial ice edge community (T0) and to subsampled volume from treatments at T1, T3 and T7. After incubations (∼ 6 hours), the microbial communities (>0.7 μm) were collected on GF/F filters (Whatman; combusted at 450°C for 2 hours), and stored frozen. Isotopic enrichments of ^15^N and ^13^C were measured on a Europa 20/20 isotope ratio mass spectrometer, and absolute uptake rates (μM h^-1^) were calculated according to Dugdale and Wilkerson 1986. Complete methods and uptake rate measurements from all time points (Fig. S1) can be found in the SI.

## Data Availability

The data reported in this paper have been deposited in the NCBI sequence read archive (BioProject accession no. PRJNA637767; BioSample accession nos. SAMN15154229 - SAMN15154256). Contigs, assembled ORFs, and MCL cluster abundance and differential expression analysis results are available at https://datadryad.org/stash/share/o4oDU_xXkHACRHkrxsKuKlDcGXjlIpdTn9I7CzaUb1Q

## Acknowledgements

We are grateful to Antarctic Support Contractors, especially Ned Corkran, for facilitating fieldwork. We thank Jeff Hoffman, Zhi Zhu, and Quinn Roberts for assistance in the field and Hong Zheng for her work in the laboratory. This study was funded by National Science Foundation (NSF) Antarctic Sciences Awards 1103503 (to E.M.B.), 0732822 and 1043671 (to A.E.A.), 1043748 (to D.A.H.), 1043635 (to D. A. B.), Gordon and Betty Moore Foundation Grant GBMF3828 (to A.E.A.); NSF Ocean Sciences Award NSF-OCE-1136477 and NSF-OCE-1756884 (to A.E.A.) and 1638804 (to D.A.H.), Nova Scotia Graduate Scholarship to L.J., NSERC CGS Postgraduate scholarship and Transatlantic Ocean System Science & Technology scholarship to J.S.P.M, NSERC Discovery Grant RGPIN-2015-05009 to E.M.B., Simons Foundation Grant 504183 to E.M.B., and Canada Research Chair support to E. M. B.

## Supplementary Information

### Full Materials and Methods

#### Experimental Design

The planktonic microbial community was sampled at 3 m depth via diaphragm pump from the sea ice edge in McMurdo Sound, Antarctica on January 15^th^, 2015 (165°24.7985’E, 77°37.1370’S) from 11:00 - 12:15 using trace metal clean techniques previously described (Bertrand et al. 2015). In-situ water temperature at the time of sampling was −1 °C. The community was protected from light upon sampling using dark trash bags, stored at 0 °C until 17:00 and then split into trace-metal cleaned polycarbonate bottles (two 1.1L and one 2.7L per treatment), with and without iron supplementation at three different temperatures. Bottles were kept at −0.5 ± 0.2 °C, 3 ± 0.5 °C or 6 ± 0.5 °C at constant 65-85 uE m^-2^ sec^-1^ irradiance in indoor incubators for a total of 7 days. For iron supplmentation, 2 nM iron was added as Fe(NO_3_)_3_ from an ultrapure analytical standard solution, 1001 mg L^-1^ in 2% nitric acid. This was diluted to a working stock in pH 2.5 milli Q water with hydrochloric acid, resulting in a negligible nitrate addition to iron amended bottles.

#### Metatranscriptome Sampling and Assembly

Four separate metatranscriptome samples were taken from the initial community, one at the sea ice edge (IE; approximately 2L) and triplicates in the laboratory during bottle incubation setup (T0, approximately 2L). Subsamples of single replicates from each experimental treatment were harvested on January 16^th^ (T1) and subsamples of each replicate (n = 3 for each experimental treatment) were taken again on January 20^th^ (T5). Each was harvested onto 0.2 μm Sterivex™ filters. RNA was extracted and sequenced via paired end Illumina HiSeq. Total RNA was extracted using Trizol reagent (Thermo Fisher Scientific). Ribosomal RNA was removed with Ribo-Zero Magnetic kits (Illumina). A mixed Removal Solution was prepared from plant, bacterial, and human/mouse/rat Removal Solution at a ratio of 2:1:1. The resulting rRNA subtracted RNA was purified and subjected to amplification and cDNA synthesis, using the Ovation RNA-Seq System V2 (TECAN). One microgram of the resulting high-quality cDNA pool was fragmented to a mean length of 200 bp, and libraries were prepared using Truseq kit (Illumina) from the -repair step in the manufacturers protocol and subjected to paired-end sequencing via Illumina HiSeq.

Illumina paired reads were filtered to eliminate primer sequences and quality trimmed to Phred Q33, and rRNA identified and removed using riboPicker (Schmieder et al. 2012) (average 13.5 % rRNA). Transcript contigs were assembled de novo using CLC Assembly Cell (http://www.clcbio.com) and ORFs predicted using FragGeneScan (Rho et al. 2010). Reads were mapped to ORFs using CLC (73% read mapping), and ORFs were annotated for putative function using hidden Markov models and blast-p against PhyloDB (Bertrand et al. 2015). ORFs were filtered to eliminate those with low mapping coverage (< 50 reads total over all samples), proteins with no blast hits, and no known domains. The remaining set of ORFs were assigned to chloroplast, mitochondrial or nuclear origin based on the best blast-p hit above e-value 1e^-3^ to an organism with known organellular peptide sequences (nuclear by default), and used for further comparative analysis.

Taxonomic groups of interest were defined (Fig. S2) and each ORF was assigned to a group based on best LPI taxonomy (Podell and Gaasterland 2007; Bertrand et al. 2015). Reads per kilobase mapped (RPKM) expression values for each ORF were calculated and taxon normalized using a normalization factor representing the summed taxonomic group contribution to total nuclear-assigned reads per sample. For example, the taxon-normalized expression of an ORF assigned to *Fragilariopsis* in a particular library is given as reads mapped to that ORF/ORF length/total *Fragilariopsis* nuclear assigned reads in that library. ORFs were clustered into orthologs and protein families using MCL (Enright 2002). MCL clustering was run in label mode (parameter -abc), with the default inflation setting 2.0 (parameter -I), on ratios of best blast-p bitscore to self-hit bitscore using BLASTALL (e-value: 1e^-3^). Group normalized cluster (average/total) RPKM expression values were calculated by pooling the taxon-normalized expression values for each group within a cluster. These taxon-normalized RPKM values were used to examine gene and cluster abundance. Cluster annotations were aggregated by annotation type (Kegg, KO, KOG, KOG class, Pfam, TIGRfam, EC, GO) and a single annotation chosen to represent each cluster based on the lowest Fisher’s exact test p-value (fisher.test in R) given the 2-way contingency table for each annotation coverage of each cluster.

#### ISIP1 taxonomic re-assignment

*Pseudo-nitzschia* and *Fragilariopsis* ISIP1 genes could not be differentiated using blast-p. Instead, nucleotide ISIP1 sequences were compared to a reference database of ISIP1 genes from *F. cylindrus, F. kergulensis, P. granii, P. heimii, P. multistriata* and *P. fraudulenta* using blast-n. ISIP1 sequences were assigned to *Fragilariopsis* or *Pseudo-nitzschia* based on lowest e-value score. These sequences were then manually placed in clusters and their abundance was normalized to each taxon as previously described.

#### Query for Domoic Acid Biosynthesis (DAB) Genes

BioEdit v7.2 was used to conduct a local blast-p search for DAB genes in our data using Blosum62 similarity matrix. Amino acid query sequences from *Pseudo-nitzschia multiseries* DAB-A (GenBank: AYD91073.1), DAB-B (GenBank: AYD91072.1), DAB-C (GenBank: AYD91075.1) and DAB-D (GenBank: AYD91074.1) (Brunson et al. 2018) were used.

#### LHC assignments

All *Fragilariopsis* and *Pseudo-nitzschia* ORFs annotated as ‘chlorophyll binding’ or ‘light harvesting proteins’ were selected for further inspection. For *Fragilariopsis* LHCs, a blast-p was performed against *F. cylindrus* CCMP1102 and annotations were retrieved for the top blast hit for each amino acid sequence. *Pseudo-nitzschia* LHC sequences were identified by comparison with those collected during the annotation of the *Pseudo-nitzschia multiseries* CLN-47 genome. The protein sequences of both diatoms were assigned to the Lhcf, Lhcr, Lhcx or Lhcz groups following a previous diatom LHC classification based on maximum-likelihood phylogenetic trees and published as Supp. Information 11 and Supp. Fig. 20 of Mock et al. (2017) and in Hippmann et al. (2017). The dominant Lhcf clade was further subdivided into Lhcf_I, Lhcf_II (diatom-specific), and Lhcf_III groups. PID numbers and the clusters to which they were assigned (see above) can be found in Table S3.

#### Plastocyanin Tree

Previously identified plastocyanin sequences were retrieved from MMETSP (*Fragilariopsis kerguelensis*_0735, *Pseudo-nitzschia heimii*_1423, *Coscinodiscus wailesii*_1066), JGI (*Fragilariopsis cylindrus*_272258), NCBI (*Thalassiosira oceanica*_EJK71623.1) and Cohen et al. 2018 (*Pseudo-nitzschia granii*), and aligned with plastocyanin sequences from *Pseudo-nitzschia* and *Fragilariopsis* in our dataset using Clustal Omega in SeaView v5.0 (Fig. S8). A maximum-likelihood phylogeny was then estimated using PhyML with LG model and 100 bootstrap iterations in SeaView v5.0, and the resulting tree was edited using FigTree v1.4.4.

#### Nutrient Measurements

Samples from initial (T0) and subsequent time points (T1, T3, T5, and T7) were collected, passed through a GF/F filter (Whatman; 0.7 μm nominal pore size; combusted at 450 °C for 2 hours), and filtrate was stored frozen (−40 °C) until further analysis. A Lachat QuickChem 8500 autoanalyzer was used to measure duplicate concentrations of dissolved nitrate, phosphate and silicate (detection limit 0.03 μmol N L^-1^, 0.03 μmol P L^-1^ and 0.05 μmol Si L^-1^; Parsons et al. 1984). Samples for ammonium, collected on T0 and T7, were measured in triplicate on a Shimadzu UV-1601 spectrophotometer using the manual phenol-hypochlorite method (detection limit 0.05 μmol N L^-1^; Koroleff 1983).

#### Uptake Measurements

Nitrate and bicarbonate uptakes were assessed using the initial community (T0) collected from the ice edge and during T1, T3, and T7 of the experiment (Fig. 1 and Fig. S1). Uptake rates were measured using ^15^N and ^13^C stable isotope tracer techniques, and substrates used included ^15^N-labeled potassium nitrate (K^15^NO_3_^-^; 98%) and ^13^C-labeled sodium bicarbonate (NaH^13^CO_3_^-^; 99%; both substrates came from Cambridge Isotope Laboratories, Andover, MA). Uptake experiments at T0 were done in triplicate using 1 L polyethylene terephthalate glycol-modified bottles. At T1, T3, and T7 a single replicate of each treatment was subsampled, and uptake experiments were done in duplicate in 230 mL polycarbonate conical bottles (all bottles were acid washed with 10% hydrochloric acid and thoroughly rinsed with ultrapure water). After tracer level additions (less than 10% of background concentrations) of ^15^N and ^13^C-labled substrates were made, bottles were returned to their respective incubators for approximately 6 hours. Incubations were terminated by filtering microbial communities (> 0.7 μm) onto combusted (450 °C for 4 hours) Whatman GF/F filters. During T0 incubations, two microbial size fractions (0.7 – 5.0 μm collected on GF/F filters and > 5.0 μm collected on Sterlitech silver membrane filters) were added together to represent the > 0.7 μm microbial community. Filters were kept fozen (−40 °C) inside 1 mL cryo vials until particulate nitrogen and carbon concentrations and isotopic enrichment of ^15^N and ^13^C were measured on a Europa 20/20 isotope ratio mass spectrometer. Absolute uptake rates for ^15^N-labeled nitrate and ^13^C-labeled bicarbonate were calculated according to Dugdale and Wilkerson (1986) and Hama et al. (1983) respectively. Nitrate uptake rates were not corrected for isotope dilution because concentrations of nitrate were greater than 5.5 μmol N L^-1^ at all time points, and isotope dilution is generally negligible when concentrations are high (Baer et al. 2017).

#### Cell Counts

Phytoplankton cell count samples from the initial (T0) and final days (T7) of the experiment were preserved with 1% glutaraldehyde, stored refrigerated in the dark and later enumerated in the lab on a Sedgwick Rafter counting chamber using an inverted compound light microscope (Accu-Scope 3032), as in (Tatters et al. 2018). All plankton taxa were identified to the lowest taxonomic level possible according to (Tomas 1997) and (Scott and Marchant 2005), with special attention to the diatom genera *Fragilariopsis* and *Pseudo-nitzschia*.

#### Statistical Analysis

We analyzed differential gene expression at T5 within observed taxa to determine which genes are responsive to Fe and temperature treatments in each group. First, we normalized reads mapped to each ORF by the abundance of nuclear reads assigned to that taxonomic group in total, which controls for changes in community composition across treatments. We then used a generalized linear model with one categorical explanatory variable, with each category representing a unique experimental treatment. To examine the effect of Fe, temperature, and their interaction on gene expression, we specified model contrasts. For Fe, we tested the difference between the sum of coefficient estimates for all Fe treatments, minus the sum of coefficient estimates for all -Fe treatments. We followed a similar approach for temperature, where we tested the difference between the sum of coefficient estimates for one temperature treatment versus another temperature treatment. For both approaches, we divided these differences by the number of treatments in the sum (i.e. 3 for Fe test and 2 for each temperature test). To test for statistical significance, we used empirical Bayes quasi-likelihood F-tests (glmQLFTest in edgeR). Throughout, we used a p-value cut off of 0.05 for statistical significance.

To examine the effects of iron on gene differential expression and fold-change magnitude, -Fe treatments for all temperatures at T5 (−Fe −0.5 °C, -Fe 3 °C, -Fe 6 °C) were compared to +Fe treatments for all temperatures at T5 (+Fe −0.5 °C, +Fe 3 °C, +Fe 6 °C). For temperature effect, - 0.5 °C was compared to 3 °C at all iron conditions at T5 (+Fe and -Fe at −0.5 °C vs +Fe and -Fe at 3 °C) and −0.5 °C was compared to 6 °C at all iron conditions at T5 (+Fe and -Fe at −0.5 °C vs +Fe and -Fe at 6 °C). For a gene to be considered upregulated with temperature, it had to be upregulated in both −0.5 °C vs 3 °C and −0.5 °C vs 6 °C (same rule applied for down regulation). If a gene is upregulated in one temperature comparison and downregulated in the other, it was not considered differentially expressed. Fold-change for temperature effect was calculated from −0.5 °C (+Fe and -Fe) vs 6 °C (+Fe and -Fe).

K.O. term enrichment analysis was performed on significantly upregulated (enriched) and downregulated (depleted) ORFs separately using KEGG enrichment functions in the GOstats R package (Falcon and Gentleman 2007). A hypergeometric distribution test was used to test for significant enrichment and depletion at p < 0.05. K.O. terms associated with less than 10 ORFs were excluded from the results.

**Figure S1.**
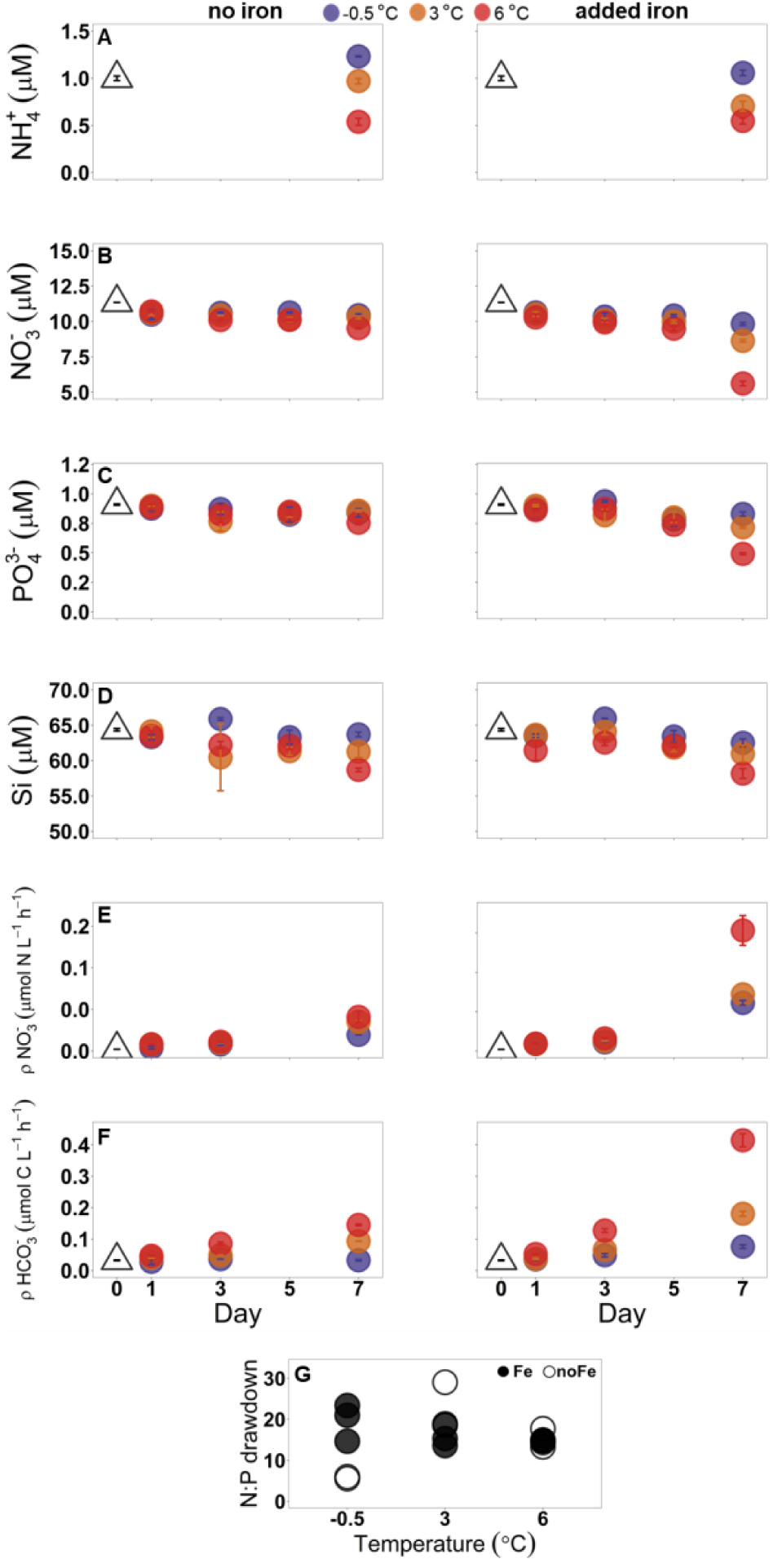
**A-D)** Dissolved ammonium, nitrate and phosphate concentrations prior to incubations (day 0), and after 1, 3, 5 and 7 days of incubation with and without added iron at - 0.5, 3, and 6 °C. Each point represents a mean value, where n = 2 on day 1, and n=3 on days 0, 3, 5, 7. In A-J, error bars represent ± 1 SD, and fall within the bounds of the symbol when not visible. **E**,**F)** Nitrate and bicarbonate uptake rates prior to incubations (day 0), and after 1, 3, and 7 days of incubation with and without added iron at −0.5, 3, and 6 °C. Each point represents a mean value, where n = 2 on days 1, 3, 7 and n = 3 on day 0. **G)** Dissolved nitrogen (nitrate + ammonium) : phosphate drawdown ratio at T7. Draw down is calculated as the difference in concentration between T7 and T0.

**Figure S2.**
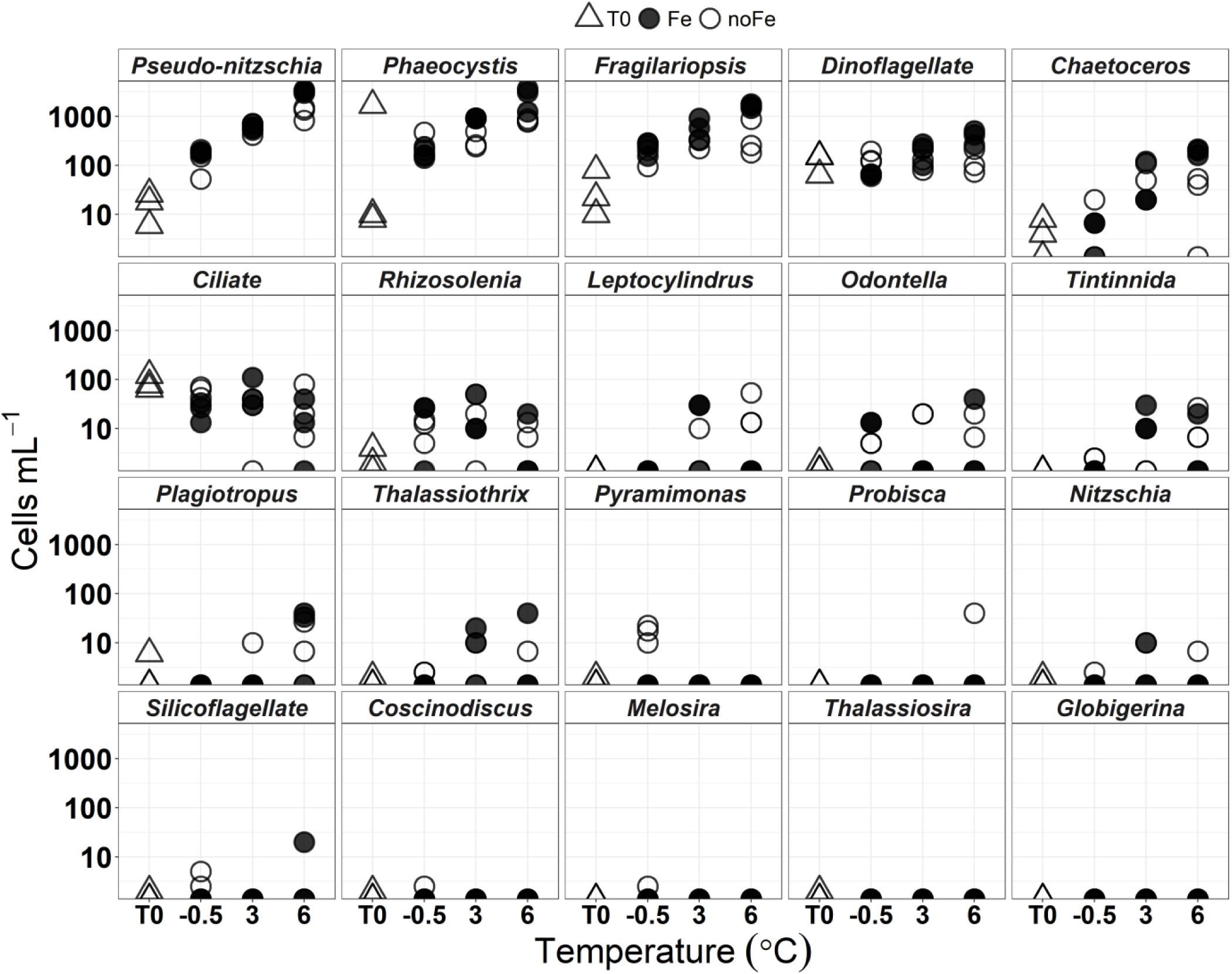
Triplicate cell count measurements of initial (T0) samples and after 7-day incubations with and without iron addition at −0.5, 3 and 6 °C. Cells from the various taxonomic groups were counted and identified using light microscopy.

**Figure S3.**
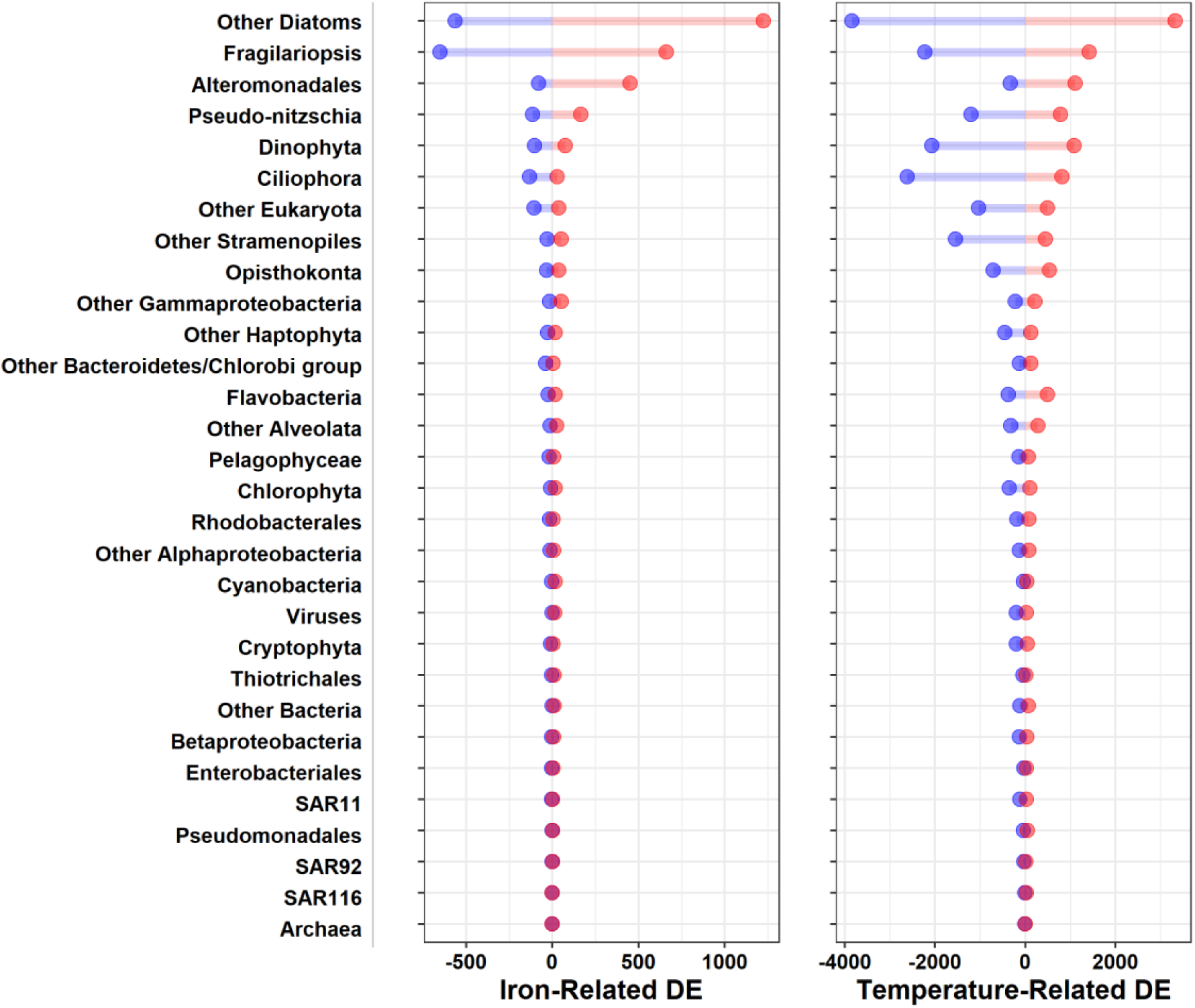
Number of significantly differentially expressed open reading frames (ORFs) belonging to the 30 taxonomic groups identified in the metatranscriptome dataset. Red = upregulation, blue = downregulation. Differential taxon-normalized expression was calculated using the quasi likelihood test (glmQLFTest) in EdgeR, and p-value cut-off of 0.05 was used for statistical significance. Iron-related DE represents differential expression patterns observed with and without iron addition at T5, temperature-related DE represents differential expression patterns observed due to warming at T5 (Methods).

**Figure S4.**
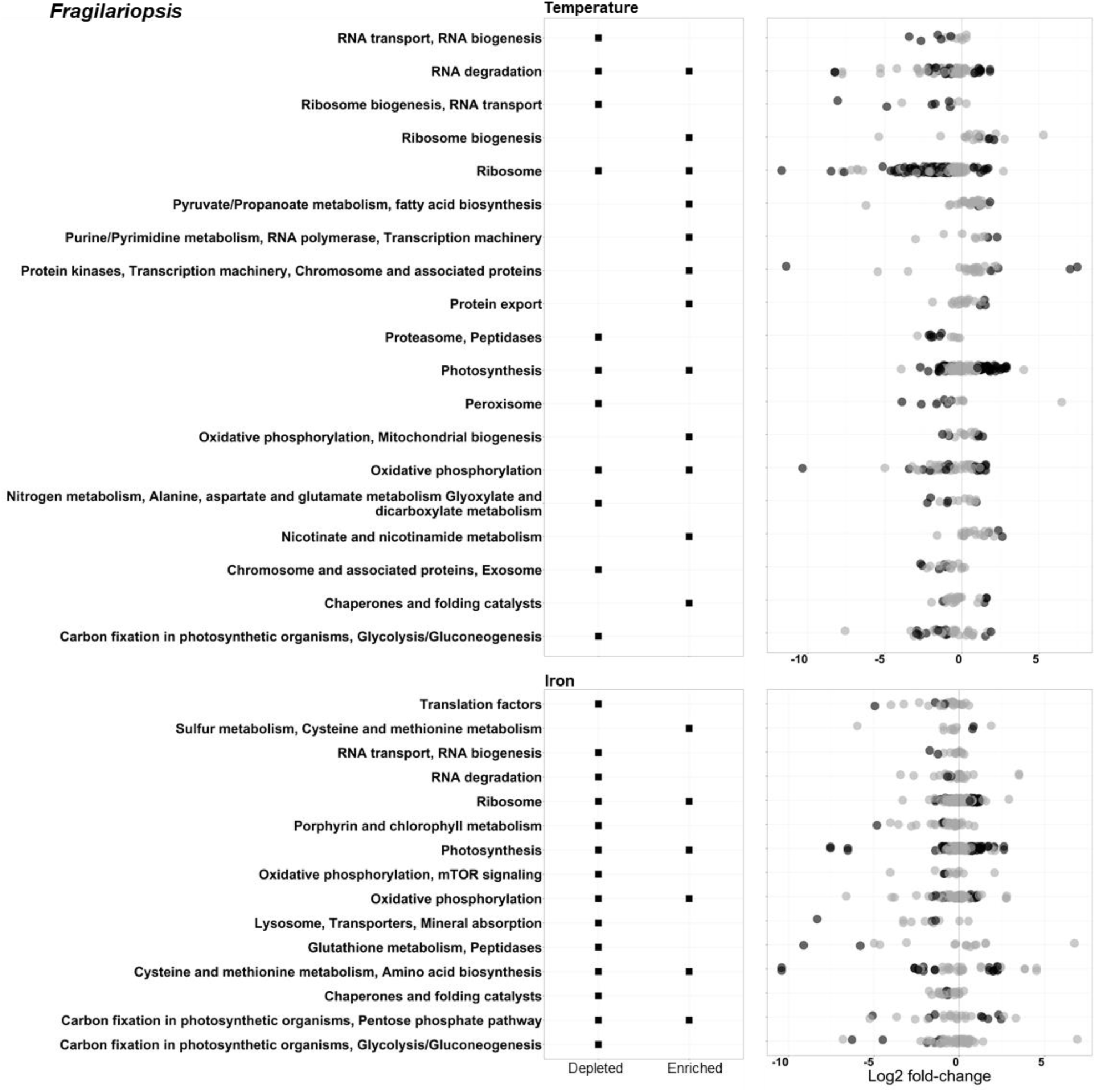
KEGG Orthology (K.O.) term enrichment analysis using all *Fragilariopsis* ORFs that were annotated with a K.O. number. Black squares correspond to ‘C’ level annotations that were significantly (p <0.05) upregulated (Enriched) and/or downregulated (Depleted) at T5 by temperature increase or iron addition. Circles correspond to the individual ORFs used in the analysis for each annotation. Black circles represent statistically significant (p <0.05) up or down regulated ORFs (positive and negative Log_2_ fold-change values, respectively). Temperature fold change was calculated using −0.5 °C vs 6 °C treatments. Iron fold-change was calculated using - Fe vs +Fe treatments at all temperatures (Methods).

**Figure S5.**
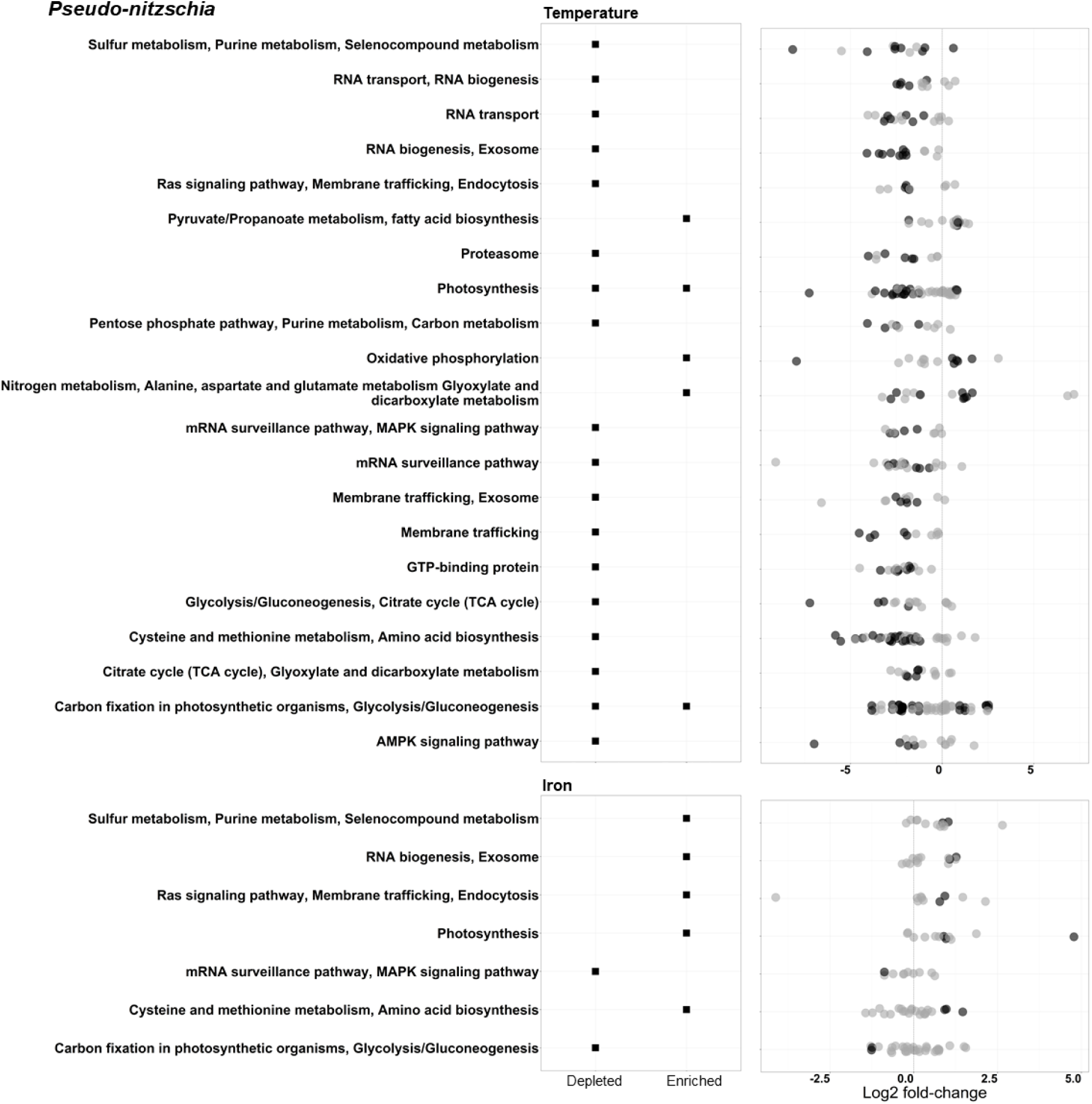
KEGG Orthology (K.O.) term enrichment analysis using all *Pseudo-nitzschia* ORFs that were annotated with a K.O. number. Black squares correspond to ‘C’ level annotations that were significantly (p <0.05) upregulated (Enriched) and/or downregulated (Depleted) at T5 by temperature increase or iron addition. Circles correspond to the individual ORFs used in the analysis for each annotation. Black circles represent statistically significant (p <0.05) up or down regulated ORFs (positive and negative Log_2_ fold-change values, respectively). Temperature fold-change was calculated using −0.5 °C vs 6 °C treatments. Iron fold-change was calculated using - Fe vs +Fe treatments at all temperatures (Methods).

**Figure S6.**
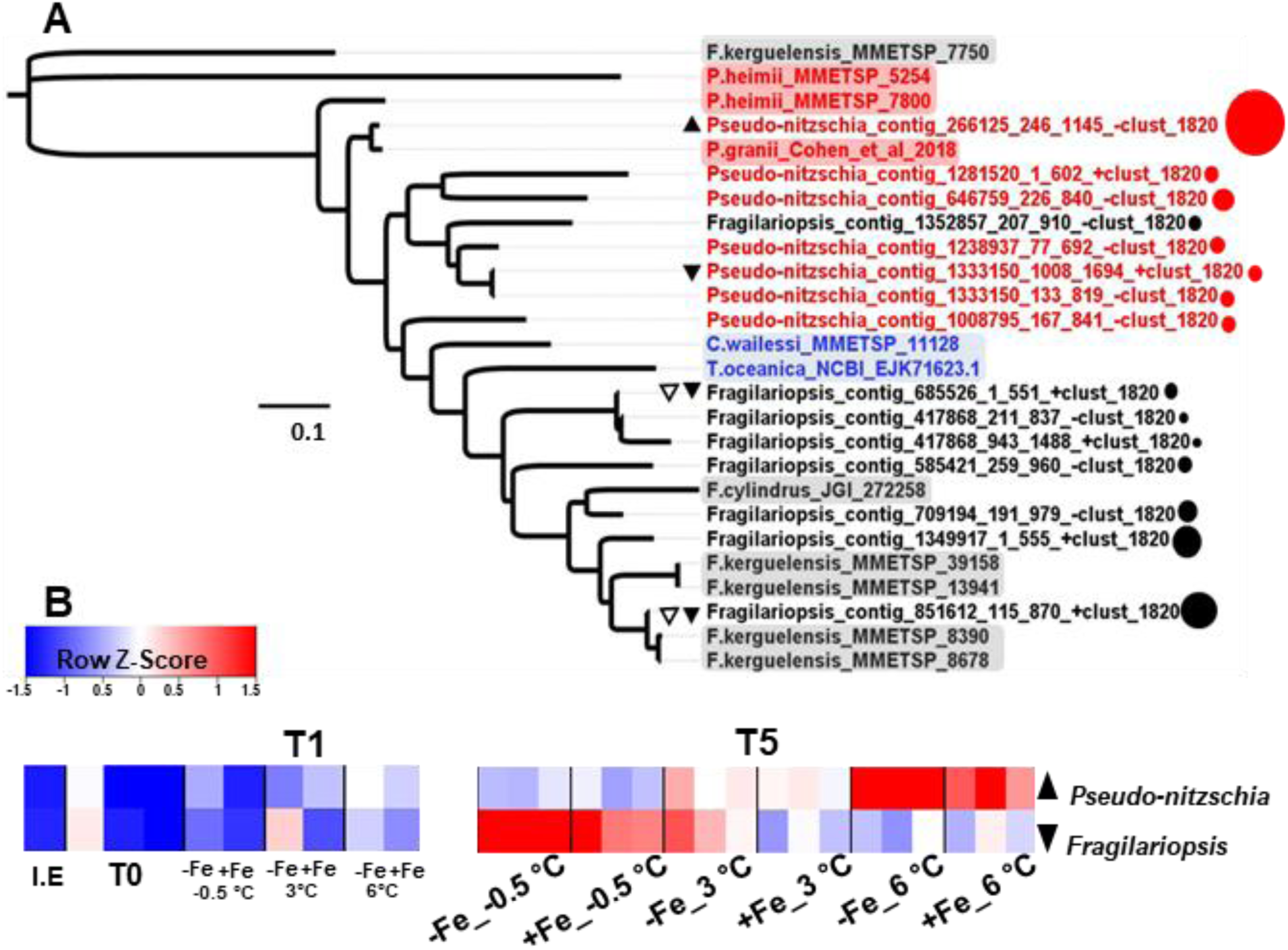
**A)** Phylogenetic tree of the plastocyanin sequences (ORFs) comprising the plastocyanin MCL cluster from both *Pseudo-nitzschia* (red) *and Fragilariopsis* (black). Non-highlighted branch tips represent ORFs in our dataset, with corresponding point size representing mean taxon-normalized ORF expression. Highlighted branch tip labels represent previously identified plastocyanin sequences retrieved from the MMETSP dataset (*Fragilariopsis kerguelensis*_0735, *Pseudo-nitzschia heimii*_1423, *Coscinodiscus wailesii*_1066), JGI (*Fragilariopsis cylindrus*_272258), NCBI (*Thalassiosira oceanica*_EJK71623.1) and Cohen et al. 2018 (*Pseudo-nitzschia granii*). **B)** Heatmaps of MCL clusters representing and plastocyanin (cluster_1820) in *Pseudo-nitzschia* and *Fragilariopsis* measured after 24 hours (T1) and 5 days (T5) of incubation under the various iron and temperature treatments. I.E represents ice edge samples, T0 represents *in-situ* samples before any incubations. Each block is one biological replicate measurement. Black-filled up/down pointing triangles represent transcripts that were significantly (glmQLFTest-EdgeR p <0.05) up or down regulated due to warming at T5.

**Figure S7.**
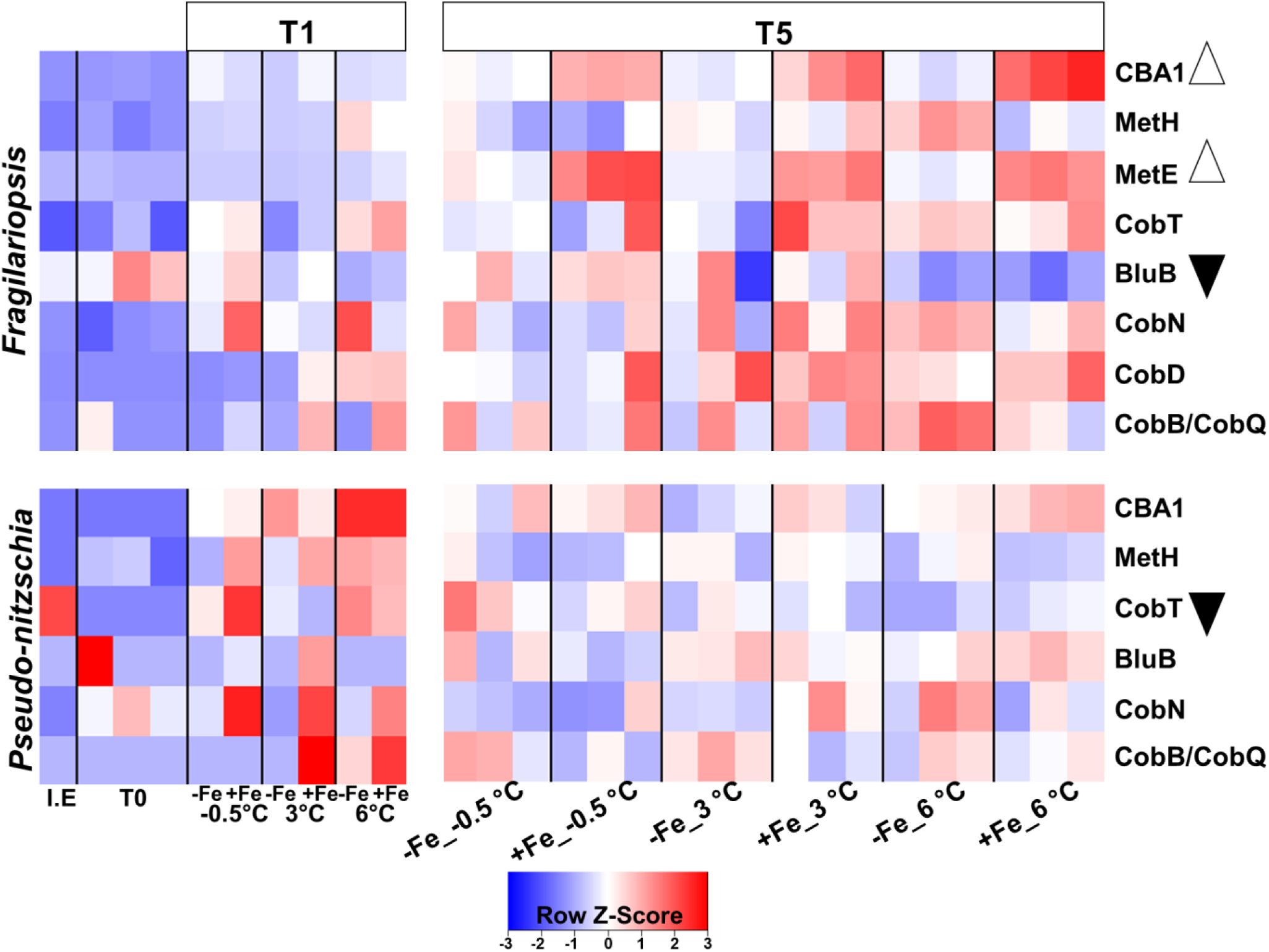
Heatmaps of MCL clusters involved in B_12_ metabolism in *Fragilariopsis* and *Pseudo-nitzschia* measured after 24 hours or 5 days of incubation under various iron and temperature treatments. I.E represents ice edge samples, T0 represents *in-situ* samples processed in the laboratory before any incubations and each block is one biological replicate measurement. CBA1: cobalamin acquisition protein 1; MetH: cobalamin-requiring methionine synthase; MetE: cobalamin-independent methionine synthase; CobT, CobN: cobaltochelatase; BluB: gene involved in DMB production; CobB/CobQ: cobyrinic acid a,c-diamide synthase/ adenosylcobyric acid synthase. Open triangles represent clusters that were significantly (glmQLFTest-EdgeR p <0.05) up regulated due to iron addition at T5. Black-filled triangles represent clusters that were significantly (glmQLFTest-EdgeR p <0.05) down regulated due to warming at T5.

**Figure S8.**
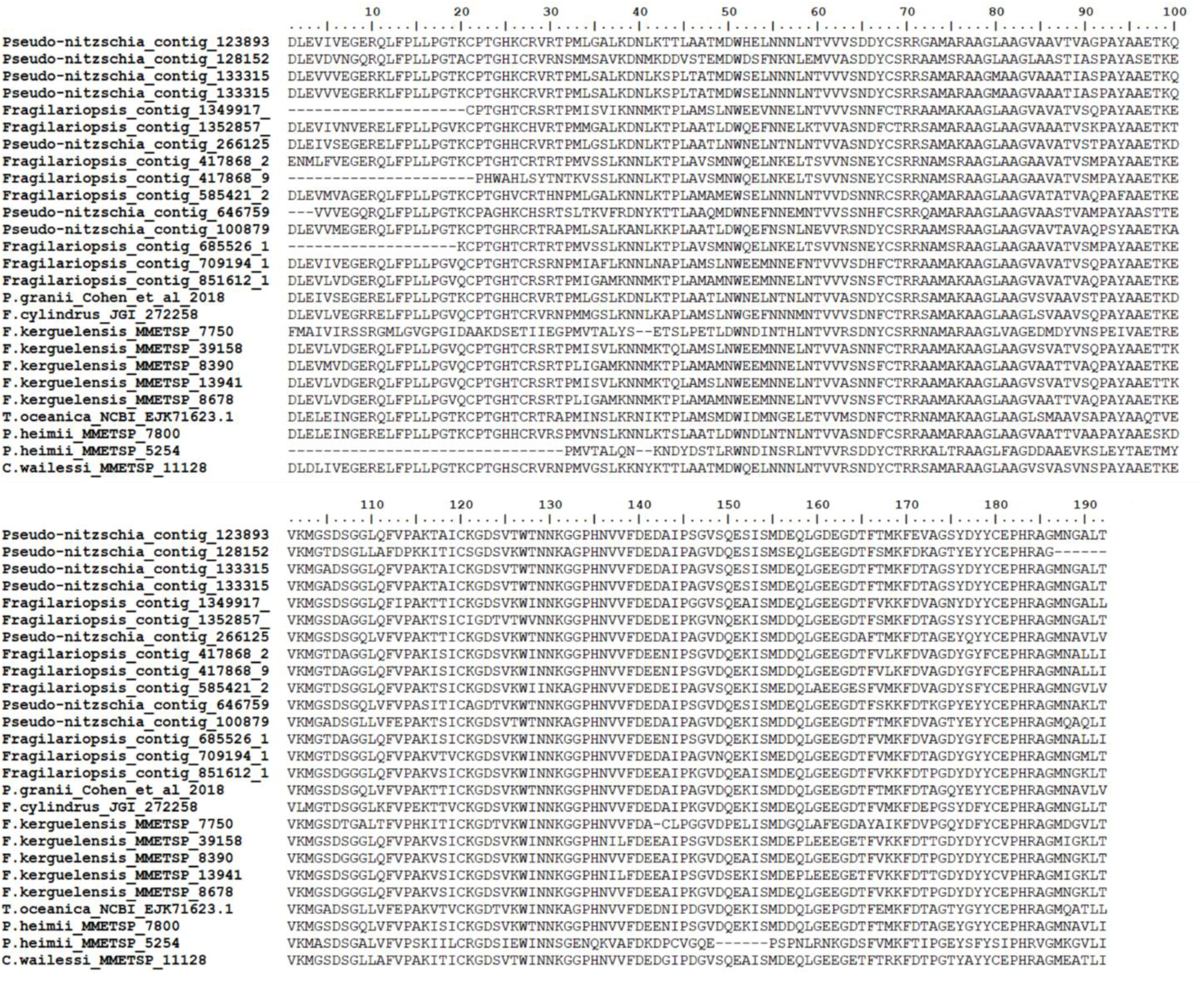
Alignment of *Pseudo-nitzchia and Fragilariopsis* plastocyanin sequences from our metatranscriptome data and previously identified plastocyanin sequences retrieved from the MMETSP dataset (*Fragilariopsis kerguelensis*_0735, *Pseudo-nitzschia heimii*_1423, *Coscinodiscus wailesii*_1066), JGI (*Fragilariopsis cylindrus*_272258), NCBI (Thalassiosira oceanica_EJK71623.1) and Cohen et al. 2018 (*Pseudo-nitzschia granii*). The alignment was conducted using Clustal Omega in SeaView v5.0. and was used to construct the maximum-likelihood phylogenetic tree for plastocyanin in Fig. S6.

**Table S1.** Blast-p search results for sequences encoding domoic acid biosynthesis proteins (DabA, B, C, D) retrieved from GenBank, against all ORFs from this study. No matches were found for DabA and DabB encoding genes and no significant eukaryotic matches were found for DabC encoding genes. Score and e-values were calculated using the Blosum62 similarity matrix. E-value 1e^-30^ was used as the cut-off for DabD results. These DabD results are further explored in Table S2.

| Query Gene | Matched Sequence (ORF ID) | Score | E-value | Taxa | Hypothesized annotation |
| --- | --- | --- | --- | --- | --- |
| **DAB-A**<br>AYD91073.1 | - | - | - | - | - |
| **DAB-B**<br>AYD91072.1 | - | - | - | - | - |
| **DAB-C**<br>AYD91075.1 | contig_254441_113_1024_+ | 63 | 6e-010 | Flavobacteria | 2OG-Fe(II) oxygenase family |
|  | contig_1174518_67_906_+ | 61 | 6e-010 | Alteromonadales | 2OG-Fe(II) oxygenase family |
|  | contig_732674_1_798_- | 56 | 9e-008 | Alteromonadales | 2OG-Fe(II) oxygenase family |
|  | contig_633551_118_1041_+ | 56 | 9e-008 | Flavobacteria | 2OG-Fe(II) oxygenase family |
|  | contig_627399_1_787_- | 54 | 5e-007 | Cyanobacteria | 2OG-Fe(II) oxygenase family |
|  | contig_1862_1_732_- | 54 | 5e-007 | Other Bacteria | 2OG-Fe(II) oxygenase family |
|  | contig_660976_1_1043_+ | 52 | 2e-006 | Other Stramenopiles | 2OG-Fe(II) oxygenase family |
|  | contig_253357_26_973_+ | 52 | 2e-006 | Flavobacteria | 2OG-Fe(II) oxygenase family |
| **DAB-D**<br>AYD91074.1 | contig_596040_24_938_- | 231 | 9e-061 | Pseudo-nitzschia | Cyt P450, CYP4/CYP19/CYP26 |
|  | contig_625510_955_1881_- | 225 | 6e-059 | Pseudo-nitzschia | Cyt P450, CYP4/CYP19/CYP26 |
|  | contig_82244_85_1023_+ | 156 | 5e-038 | Fragilariopsis | Cyt P450, CYP4/CYP19/CYP26 |
|  | contig_596040_947_1648_- | 152 | 9e-037 | Fragilariopsis | Cyt P450, CYP4X1 |
|  | contig_82244_1395_1877_+ | 145 | 7e-035 | Fragilariopsis | Cyt P450 |
|  | contig_680152_23_630_- | 132 | 6e-031 | Cyanobacteria | Cyt P450 |
|  | contig_1109498_1_752_- | 130 | 3e-030 | Other Diatom | Cyt P450 |

**Table S2.**
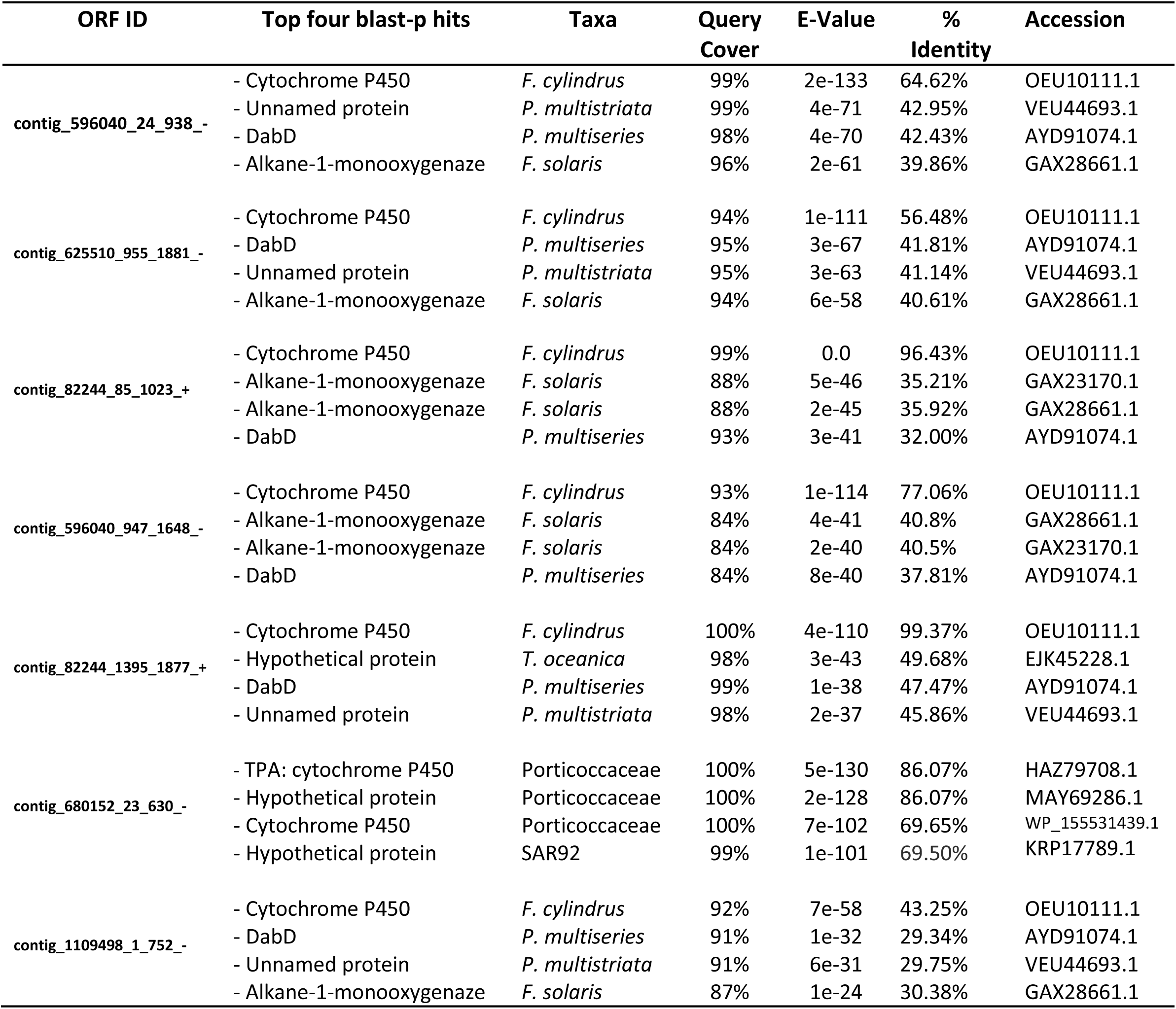
Reciprocal blast-p search results for ORFs with similarity to DabD-encoding genes (Table S1) against NCBI’s nr database.

**Table S3.**
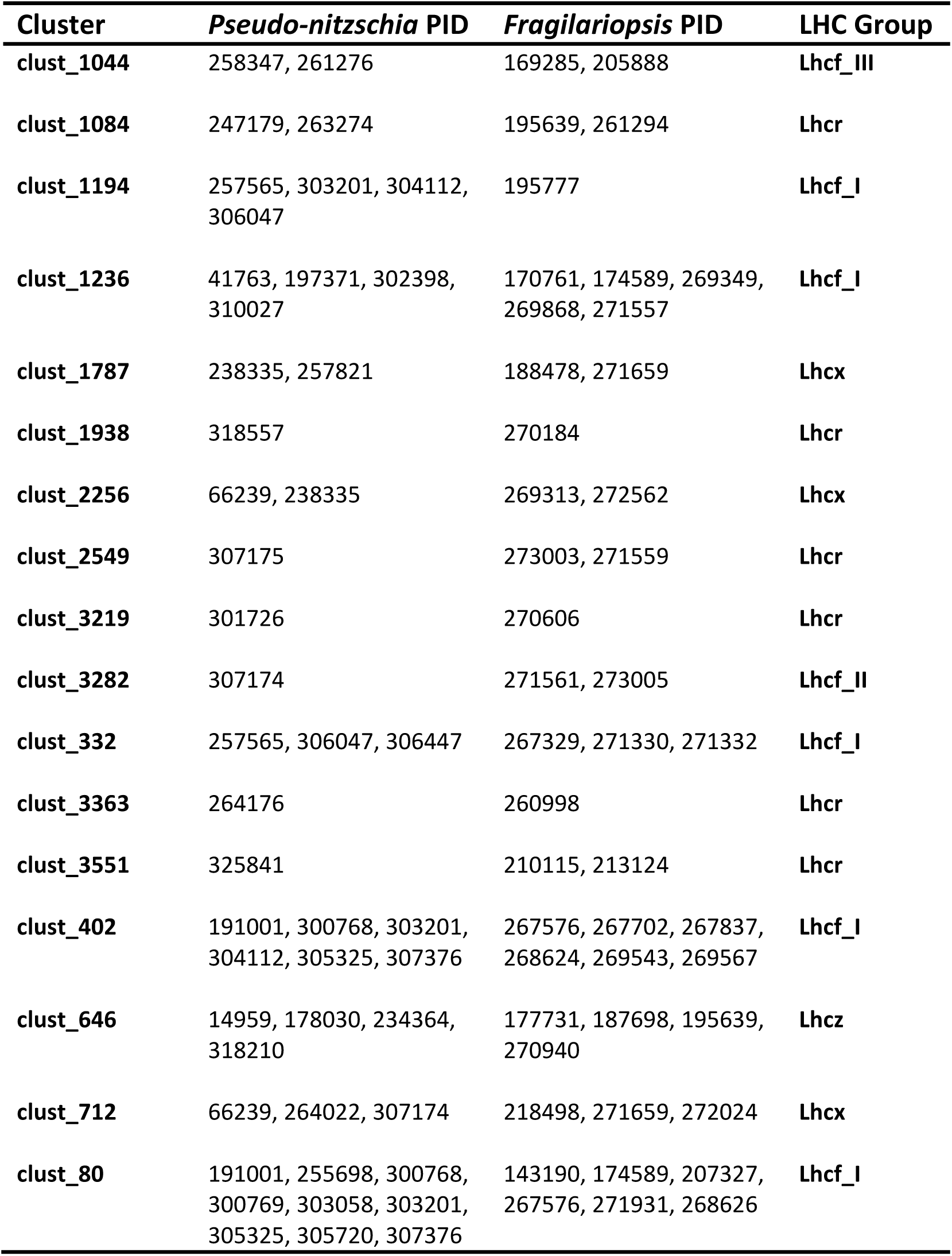
Protein ID (PID) assignments for ORFs in the various *Pseudo-nitzschia* spp. and *Fragilariopsis* spp. light harvesting complex (LHC) clusters. PIDs were assigned by performing a blast-p search against *Pseudo-nitzschia multiseries* (CLN-47) and *Fragilariopsis cylindrus* (CCMP 1102). LHC groups were assigned based on previous diatom LHC classifications in Mock et al. 2017 (Supp.Info.11) and Hippmann et al. 2017.

